# The Integrative Studies on the Functional A-to-I RNA Editing Events in Human Cancers

**DOI:** 10.1101/2022.06.05.493160

**Authors:** Sijia Wu, Zhiwei Fan, Pora Kim, Liyu Huang, Xiaobo Zhou

## Abstract

A-to-I RNA editing, constituting nearly 90% of all RNA editing events in human, has been reported to contribute to the tumorigenesis in diverse cancers. However, the comprehensive map for functional A-to-I RNA editing events in cancers is still insufficient. To fill this gap, we systematically and intensively analyzed multiple tumorigenic mechanisms of A-to-I RNA editing events in samples across 33 cancer types from The Cancer Genome Atlas. For individual candidate among ~ 1.5M quantified RNA editing events, we performed diverse types of down-stream functional annotations. Finally, we identified 24,236 potentially functional A-to-I RNA editing events, including the cases in *APOL1, IGFBP3, GluA2, BLCAP*, and *miR-589-3p*. These events showed significant results and might play crucial roles in the scenarios of tumorigenesis, due to their tumor-related editing frequencies or probable effects on altered expression profiles, protein functions, splicing patterns, and miRNA regulations of tumor genes. Our functional A-to-I RNA editing events (https://ccsm.uth.edu/CAeditome/) will help better understanding of cancer pathology from A-to-I RNA editing aspect.

## Introduction

Adenosine-to-inosine (A-to-I) RNA editing is the most common RNA editing type in humans, constituting nearly 90% of all RNA editing events. Recently, increasing evidence has revealed a significant contribution of RNA editing to tumorigenesis through multiple mechanisms (1–3), including alteration of protein-coding capacity, generation of diverse protein isoforms, and change of cellular fate of RNA and its likelihood of being translated (3). Specifically, A-to-I RNA editing in coding sequences can result in the functional alterations of proteins that have roles in tumors. For example, an A-to-I RNA editing of *SLC22A3*, resulting in the substitution of asparagine 72 to aspartate, drives early tumor invasion and metastasis in familial esophageal cancer (4). A study of gastric cancer reported that editing at codon 241 of *PODXL* confers a loss-of-function phenotype that neutralizes the tumorigenic ability of the unedited gene (5). Also, A-to-I RNA editing can modulate splicing to generate diverse isoforms associated with cancer. In acute myeloid leukemia, an experiment in vitro provided the evidence of aberrant intron-retaining splice variant caused by the hyper-editing of *PTPN6* which is potentially involved in leukemogenesis (6). *STAT3β*, the tumor regression-associated isoform, is preferentially induced by an A-to-I RNA editing event residing in proximity to the alternatively spliced exon (7). Besides, micro RNAs (miRNAs) and the three prime untranslated regions (3’-UTRs) of mRNAs can also undergo A-to-I RNA editing, which may affect their interactions in cancer. For example, an RNA editing site in *miR-200b* has been reported to switch the functional roles of this miRNA in terms of cell migration and invasion from suppression to promotion (8). The edited mature *miR-455-5* caused the reduction of tumor growth and metastasis by promoting tumor suppressor gene *CPEB1* in melanoma (9). As shown in these examples, the systematic and intensive analyses of A-to-I RNA editing will provide critical evidence and novel therapeutic targets in human cancers.

To date, there are several pan-cancer editing landscapes covering the functional annotations of RNA editing events from the aspects of clinical associations (10–12), protein recoding (10), and miRNA regulations (8, 11). For these aspects, they either provided limited candidates for each cancer type, or included partial analyses of RNA editing. For a more comprehensive map of functional A-to-I RNA editing events, in this study, we performed a systematic and intensive bioinformatics analysis pipeline (**Fig. 1**) for all the samples across 33 cancer types from The Cancer Genome Atlas (TCGA), similar as that used in the data analyses of Alzheimer’s disease from our recent study (13). All the analyses were cooperative to point out 24,236 functional A-to-I RNA editing candidates and present their potential roles in the scenarios of tumorigenesis. From the analyses, we confirmed the possible functions of the well-known R/G editing *(GluA2*, CAediting_390714) in neurological and brain tumors, expanded the roles of *BLCAP* Q/R editing (CAediting_1426931) in carcinogenesis promotion in pan-cancers, and re-addressed the tumorigenic control potential of edited *miR-589-3p* (CAediting_524911) through dysregulations of tumor genes. In addition, we also identified two another novel and promising functional RNA editing events. One case (CAediting_1478179) was up-edited in diverse cancers and may confer its pathological function through the intervention in miRNA regulation on the tumor gene of *APOL1*. Another event (CAediting_543208) occurred only in tumor samples for multiple cancer types and may enhance the ability of *IGFBP3* to inhibit tumor cell growth. All these discoveries are available at https://ccsm.uth.edu/CAeditome/. This database provides novel knowledge of tumorigenesis and list potential targets for cancer and drug research communities.

**Figure 1.**
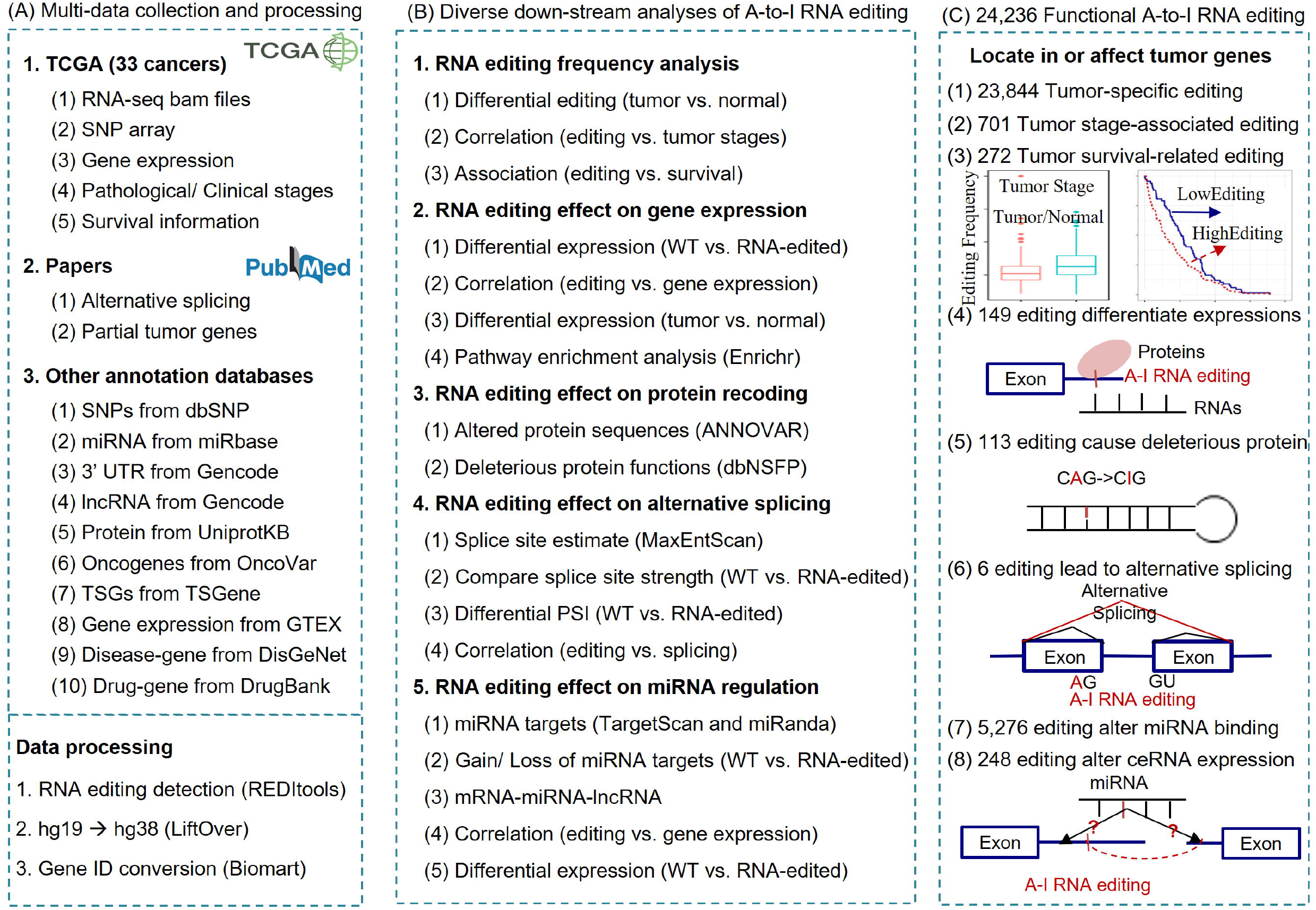
The flowchart to identify functional A-to-I RNA editing events in cancers. It describes (A) the collection and pre-processing of multi-omics data across 33 cancer types, (B) diverse down-stream analyses of A-to-I RNA editing events, and (C) potentially functional A-to-I RNA editing events related to tumorigenesis.

## Results

### 23,904 RNA editing candidates were abnormally edited in cancer

The changes of editing frequencies in tumors, along with diverse stages of tumor pathology, and across different tumor survival statuses can reveal aberrant RNA editing events which may be responsible for tumor occurrence, progression, and poor survival. In this work, after comparing editing frequencies between tumor samples and controls, we identified 23,844 RNA editing events in 869 tumor genes showing tumor-specific frequencies (**Fig. 2**A-F, Table S1, Fig. S1-4). Next, through the correlation studies of editing frequencies with tumor stages, we found 701 RNA editing events in 158 tumor genes which were significantly associated with tumor progression (Fig. 2G-I, Table S1, Fig. S5-6). Then the survival analysis discovered 272 RNA editing events in 99 tumor genes which might affect the survival risks of cancer patients (Fig. 2J-L, Table S1, Fig. S7-8). Among the 23,904 functional RNA editing events, we selected two candidates to show the effects of A-to-I RNA editing on tumors.

**Figure 2.**
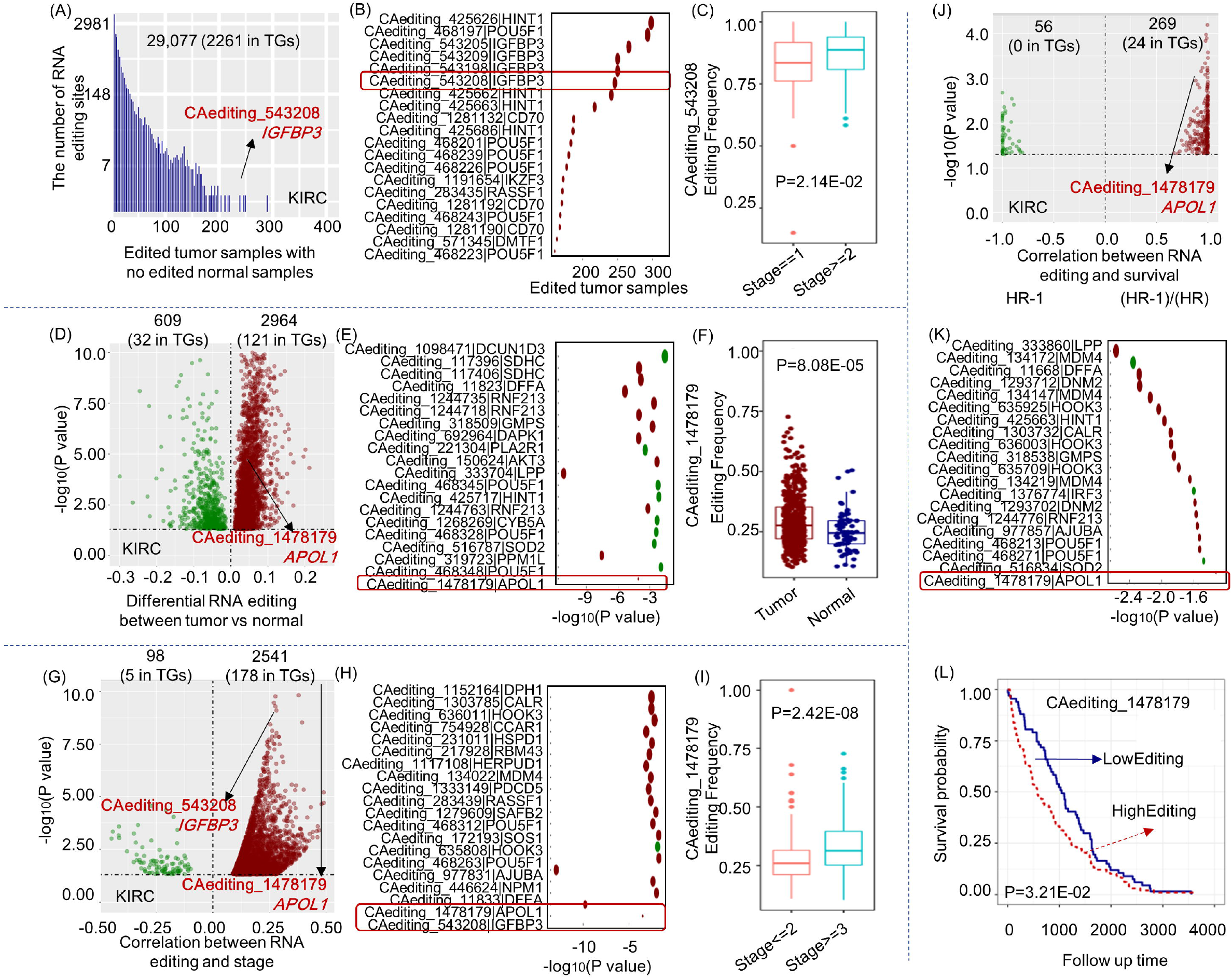
RNA editing frequencies analysis. (A-C) Kidney renal clear cell carcinoma (KIRC) specific A-to-I RNA editing events with more than five edited tumor samples and none edited normal controls. The histogram presents the distribution of this kind of RNA editing events along with the number of edited tumor samples. The bubble plot introduces a part of this kind of RNA editing events in tumor genes (TGs). One significant case of *IGFBP3* occurred only in 246 tumor samples for the KIRC cancer type, also showing higher editing frequencies in more severe KIRC tumors. (D-F) KIRC-specific A-to-I RNA editing events showing differential editing frequencies in KIRC tumors compared to controls. The volcano plot presents the differences of editing frequencies between tumor samples and controls. The bubble plot introduces a part of this kind of RNA editing events in tumor genes. One significant case in *APOL1* showed higher editing frequencies in KIRC tumor samples. (G-I) KIRC stage associated A-to-I RNA editing events. The volcano plot presents the correlations of editing frequencies with tumor stages. The bubble plot introduces a part of this kind of RNA editing events in tumor genes. One significant case in *APOL1* showed higher editing frequencies in more severe KIRC tumors. (J-L) KIRC survival related A-to-I RNA editing events. The volcano plot presents the correlations of editing frequencies with cancer survival. The bubble plot introduces a part of this kind of RNA editing events in tumor genes. One significant case in *APOL1* showed higher editing frequencies in the poorer survival group. The editing frequencies analysis results for other cancer types were displayed in Fig. S1-8.

One significant case is an RNA editing event in chr22: 36266650 (CAediting_1478179) of the *APOL1* gene in the cancer type of kidney renal clear cell carcinoma (KIRC). This event showed significantly higher editing frequencies in tumor samples than controls with a P-value of 8.08E-05 (Fig. 2F), was up-edited in more severe tumor samples (t-test: P = 2.42E-08, Fig. 2I; Spearman test: P = 1.71E-10 and R = 0.28), and presented a high risk for KIRC survival (KM analysis: P = 3.21E-02, Fig. 2L; COX analysis: P = 4.75E-02 and Hazard ratio = 3.42). Its existence in the 3’-UTR of *APOL1* seems to cause the up-regulation of the edited gene (t-test: P = 1.04E-14 and log2FC = 1.86, **Fig. 3**A; Pearson test: P = 7.44E-06 and R = 0.20, Fig. 3B) from the loss of original *miR-7151-3p* binding targets detected by TargetScan (14) and miRanda (15). Due to the overexpression of this gene in KIRC tumor samples compared to controls (P = 6.42E-21 and log2FC = 2.04, Fig. 3C), and its inducing role in autophagy (16), we suggest this RNA editing event as a potential biomarker of KIRC progression and survival (Fig. 3D).

**Figure 3.**
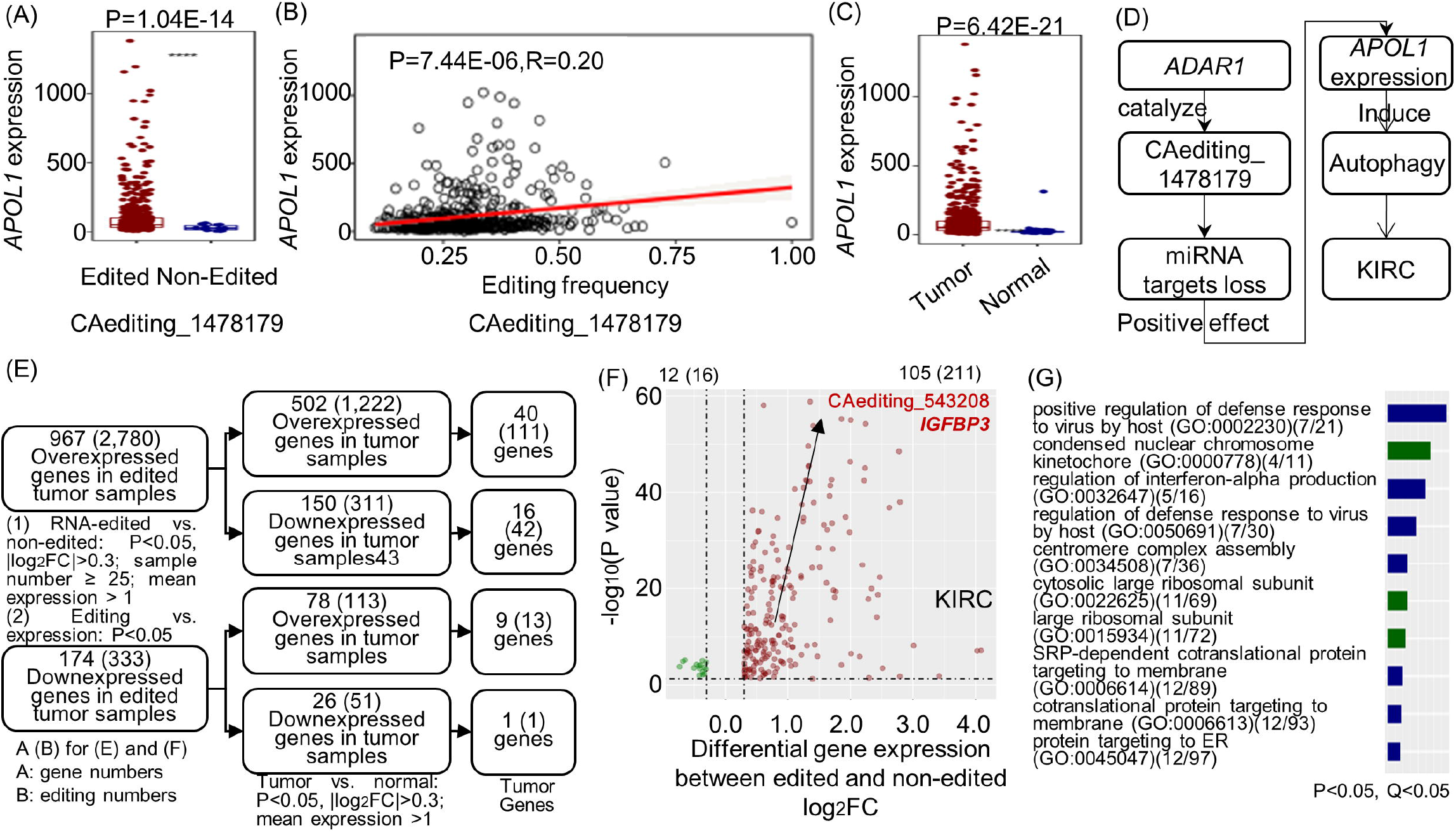
The effects of A-to-I RNA editing events on gene expressions. (A-D) One significant case in *APOL1* seems to be a potential biomarker for the KIRC cancer type. It caused the loss of original miRNA binding target to induce the over-expressions of *APOL1*, which may interfere in the autophagy function of this gene in cancer. (E) The analysis procedures for the effects of A-to-I RNA editing events on gene expressions. First, we performed differential expression genes (DEGs) analysis between RNA-edited and nonedited tumor samples, and correlation studies of gene expressions with editing frequencies to identify genes whose expressions were probably affected by A-to-I RNA editing. Then we overlapped these genes with the DEGs between tumors and controls, to focus on RNA editing effects on the aberrantly expressed genes in cancer, especially the tumor genes. (F). The overlapped DEGs in the KIRC cancer type. (G) The overlapped DEGs were enriched in the immune and replication related functions and processes. The RNA editing effects on gene expressions in other cancer types were presented in Fig. S10-13.

Another editing candidate locates in chr7: 45916046 (CAediting_543208) of the *IGFBP3* gene as shown in Fig. S9. For the KIRC cancer type, this event occurred only in tumor samples (246/535 vs. 0/72) and was up-edited in the samples with higher stages (t-test: P = 2.14E-02, Fig. 2C; Spearman test: P = 3.04E-04 and R = 0.23). Moreover, it seems to be linked with the overexpression of *IGFBP3* (t-test: P = 2.18E-09 and log2FC = 0.61; Pearson test: P = 2.06E-02 and R = 0.15). Since this gene was up-regulated in tumor samples (P = 1.81E-90 and log2FC = 3.47), and appeared to act in an autocrine action to suppress tumor cell growth (17, 18), we may carefully suggest the role of this RNA editing event in enhancing the protective functions of *IGFBP3* against cancer progression.

### 149 RNA editing candidates are potential factors to affect tumor gene expressions

From the analyses above, we found that the frequencies of several RNA editing events were significantly associated with the expressions of their host genes to be involved in tumors. For systematical analysis of the potential contributions of these RNA editing events to the expression levels of the edited genes, we performed differentially expressed gene (DEG) analysis between RNA-edited and non-edited tumor samples, and correlation studies of gene expressions with corresponding editing frequencies (Table S1, Fig. S10-11). The results were combined with the DEGs between tumor samples and controls for following enrichment analysis, to understand the probably involved pathways and biological functions of these RNA editing events in cancer.

As shown in Fig. 3E-F, we first discovered 2780 and 333 RNA editing events that would cause the up- and down-regulations of 967 and 174 edited genes respectively. Of them, 651 genes were also abnormally expressed in tumor samples compared to controls, including 55 tumor genes possibly affected by 149 A-to-I RNA editing events. Combining the potential promotion or inhibition roles of these edited genes in cancer, we could infer the possible functions of these RNA editing events related to tumorigenesis, such as CAediting_1478179 of *APOL1* and CAediting_543208 of *IGFBP3* mentioned above.

Besides, Fig. S12 presents a well-known R/G editing case in chr4:157360142 position (CAediting_390714) of the *GluA2* (syn. *GRIA2*) in the cancer type of pheochromocytoma and paraganglioma (PCPG). Its editing frequencies were positively associated with the expressions of its host gene (t-test: P = 5.04E-13 and log2FC = 1.02; Pearson test: P = 1.34E-02 and R = 0.20). Due to the over-expressions of this gene in tumors (P = 4.64E-23 and log2FC = 5.11), and its possible roles in proliferation stimulation, apoptosis resistance, migration, and invasion in cancer cell lines (19), this editing event may be a pathological biomarker for the PCPG cancer type, which was also supported by 146 edited PCPG tumor samples and none edited normal samples.

The following enrichment analysis for these 651 edited DEGs revealed immune and replication related biological functions and processes (Fig. 3G). Specifically, activation of innate and adaptive immune response through the regulation of viral defense and interferon-alpha production is benefit for cancer immunotherapy (20, 21). The replication processes related to ribosome, endoplasmic reticulum, kinetochore, and so on are important for tumor proliferation and cancer risks (22–24). Moreover, for each cancer type, the enrichment analysis also discovered some tumor-related KEGG pathways as shown in Fig. S13 and Table S2. For example, targeting apoptosis is a promising therapy to eliminate cancer cells (25), antigen processing and presentation pathway (APP) is the cellular mechanism that determines direct interactions between cancer cells and adaptive immune system (26), and sphingolipids metabolic network provides regulatory nodes for controlling tumor growth and proliferation in response to cellular stress (27). Then the DEG-associated RNA editing events probably affect these pathways or processes to be involved in cancer.

### 113 RNA editing candidates may reshape their protein functions in tumorigenesis

RNA editing events in coding regions can alter amino acid sequences and have a chance to affect protein functions. To study this, we first selected A-to-I RNA editing sites in protein-coding sequences for the identification of 3785 non-synonymous and 121 stop-loss editing events (**Fig. 4**A). Out of these, 1128 RNA editing sites were recognized to have impacts on the biological functions of 491 proteins by at least one of the annotation tools such as SIFT, Polyphen2, and PROVEAN. Among them, 113 A-to-I RNA editing events may reshape the functions of 52 tumor-related proteins (Fig. 4B).

**Figure 4.**
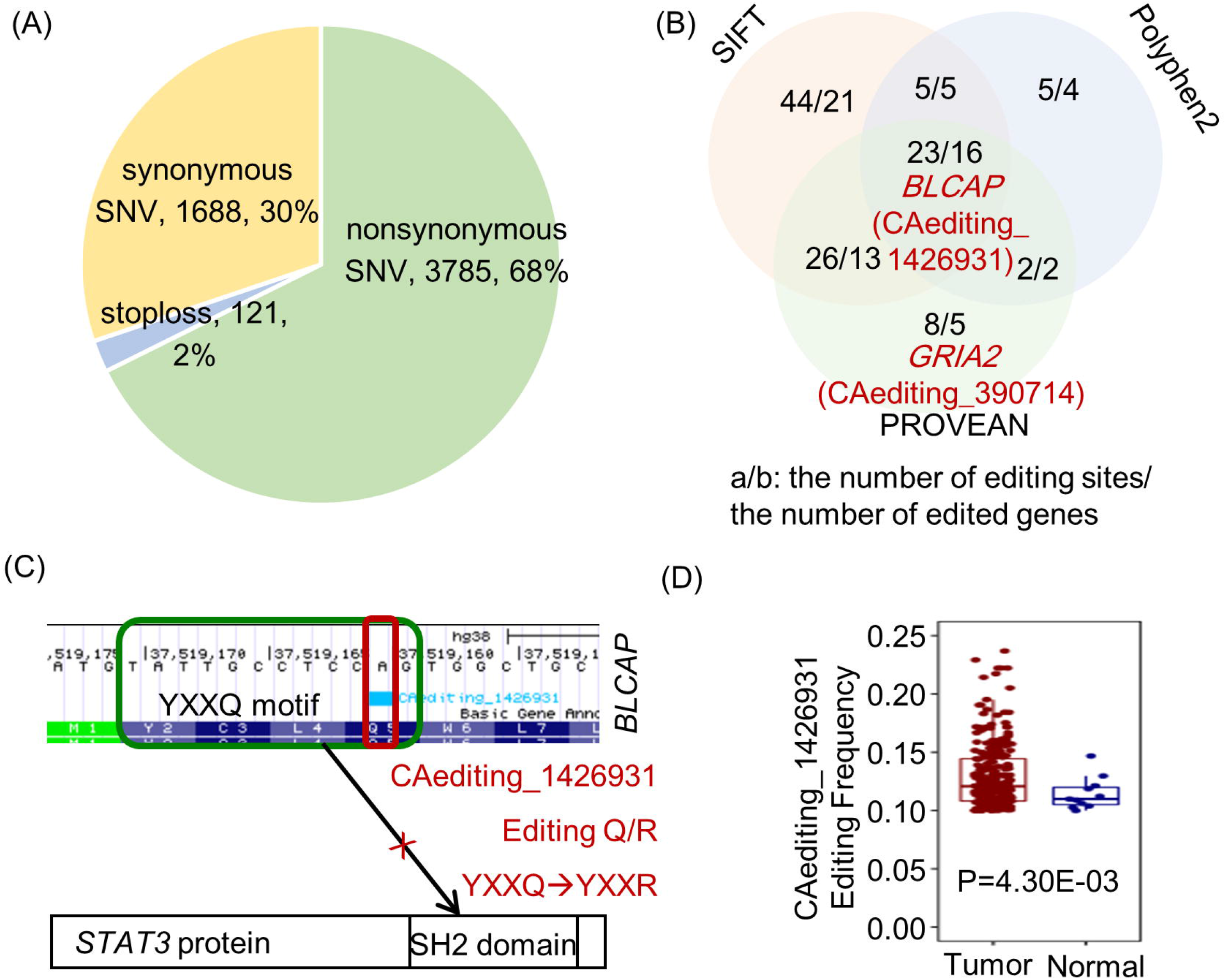
The effects of A-to-I RNA editing events on protein recoding. (A) There are 3785 non-synonymous and 121 stop-loss editing events causing the changes of amino acid sequences. (B) 113 RNA editing events conferred their deleterious effects on 52 tumor-related proteins assessed by SIFT, Polyphen2, and PROVEAN. (C) The Q/R editing in the key YXXQ motif of *BLCAP* protein reverses the inhibition ability of *BLCAP* to *STAT3*, potentially facilitating the cancer-initiating and progressing events. (D) This hypothesis was supported by its up-editing frequencies in breast invasive carcinoma.

One RNA editing candidate in the position of chr20: 37519161 (CAediting_1426931) leads to the Q/R changes of key YXXQ motif in the *BLCAP* protein (Fig. 4C). This editing event reverses the inhibition ability of *BLCAP* to *STAT3*, facilitating the cancer-initiating and progressing events (28, 29). Its roles in carcinogenesis promotion were also supported by its abnormal editing cases in pan-cancers, such as the up-editing frequencies in the cancer types of breast invasive carcinoma (BRCA, P = 4.30E-03, Fig. 4D) and KIRC (P = 4.33E-02), positive associations with tumor stages for bladder urothelial carcinoma (BLCA, P = 3.03E-02, R = 0.29), and mere occurrence in tumor samples of BLCA (55/411 vs. 0/19), colon adenocarcinoma (COAD, 176/471 vs. 0/41), head and neck squamous cell carcinoma (HNSC, 9/501 vs. 0/44), cholangiocarcinoma (CHOL, 10/36 vs. 0/9), and rectum adenocarcinoma (READ, 70/167 vs. 0/10).

Another well-known R/G editing event (CAediting_390714) in *GluA2* protein mediates the fast excitatory synaptic transmission (13, 30, 31) and may affect the functions of *GluA2* in tumor cell growth, migration, and invasion (10, 19, 32). Its roles in cancer were also supported by its differential editing frequencies in glioblastoma multiforme (GBM, P = 1.78E-02, Fig. S3). Therefore, the other 111 RNA editing events may also be possibly involved in cancer through modifying the functions of tumor-related proteins, which deserve to be studied further.

### 6 RNA editing candidates are probable regulators of alternative splicing in tumor genes

RNA editing sites in the alternatively spliced exon regions can differentiate splice site strength and eventually affect the selection of splicing positions. To study this, we focused on the editing sites locating around the exon junction boundaries for all the 33 cancer types. In total, we identified 3600 RNA editing sites in the 3’-acceptor splice regions (3’-ss) and 1779 RNA editing sites in the 5’-donor splice regions (5’-ss). They present diverse impacts on the splice site strength of 1957 genes, due to their different locations in the splicing sequences (**Fig. 5**A). Among these editing events, 79 cases have verified their effects on alternative splicing (Fig. 5B), through the differential percent spliced in (PSI) values in RNA-edited samples and significant correlations of PSI values with editing frequencies. Out of them, 6 A- to-I RNA editing events may have an opportunity to involve in autophagy reduction (16), tumor growth (33), cell proliferation (19, 32, 34), cancer metastasis (19, 32, 34), toxicity mediation (35), and so on, since they altered the splicing patterns of tumor genes.

**Figure 5.**
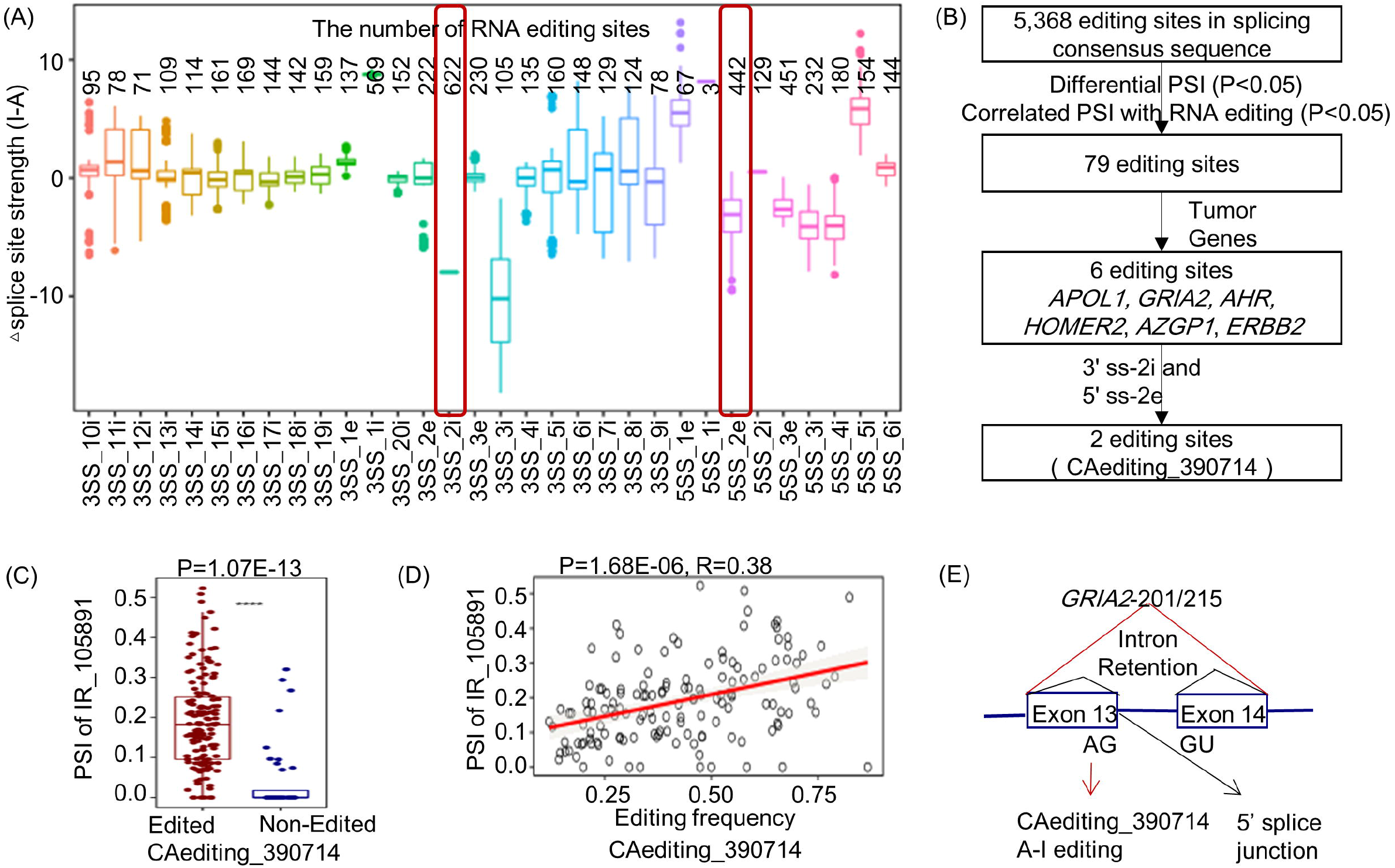
The effects of A-to-I RNA editing events on alternative splicing. (A) The distribution of altered splice site strength caused by RNA editing events in the different positions of splicing regions. (B) The analysis procedures for the effects of A-to-I RNA editing events on alternative splicing. (C-E) The R/G editing in *GluA2* altered the canonical splicing pattern of AG-GU to induce the intron retention for the isoforms of tumor-related *GluA2* in pheochromocytoma and paraganglioma.

One case of them is the well-known R/G editing (CAediting_390714) in *GluA2*. It altered the canonical splicing pattern of AG-GU and caused a reduction of the 5’-donor splice site strength (5.37 - 8.23 = −2.86) to potentially induce the intron retention (157360143:157361009) for the isoforms of *GluA2* in the PCPG cancer type (Fig. 5C-E). Combined all the analyses above, this editing event may play important roles in neurological or brain cancers through altering the excitatory synaptic transmission by its contributions to protein function changes, alternative splicing, and dysregulation of *GluA2* (Fig. S12).

### 248 RNA editing candidates likely intervened in miRNA regulations on tumor genes

RNA editing sites in the binding targets of miRNAs or their seed regions would alter miRNA-RNA interactions and possibly affect the expressions of miRNA-regulated genes. Through miRNA targets prediction for wild-type and RNA-edited transcripts, we identified 96,278 RNA editing sites in 7930 transcripts, which were presumed to potentially create 402,284 new miRNA targets and eliminate 291,685 original ones (**Fig. 6**A). These altered miRNA-targets interactions further conferred their effects on the expressions of 14,107 regulated genes (Table S1) other than the edited genes. Taking into account the functions of all these genes in tumor, we eventually selected 248 functional A-to-I RNA editing candidates, which may likely intervene in miRNA regulations on 436 tumor genes to play their roles in tumorigenesis (Fig. 6B).

**Figure 6.**
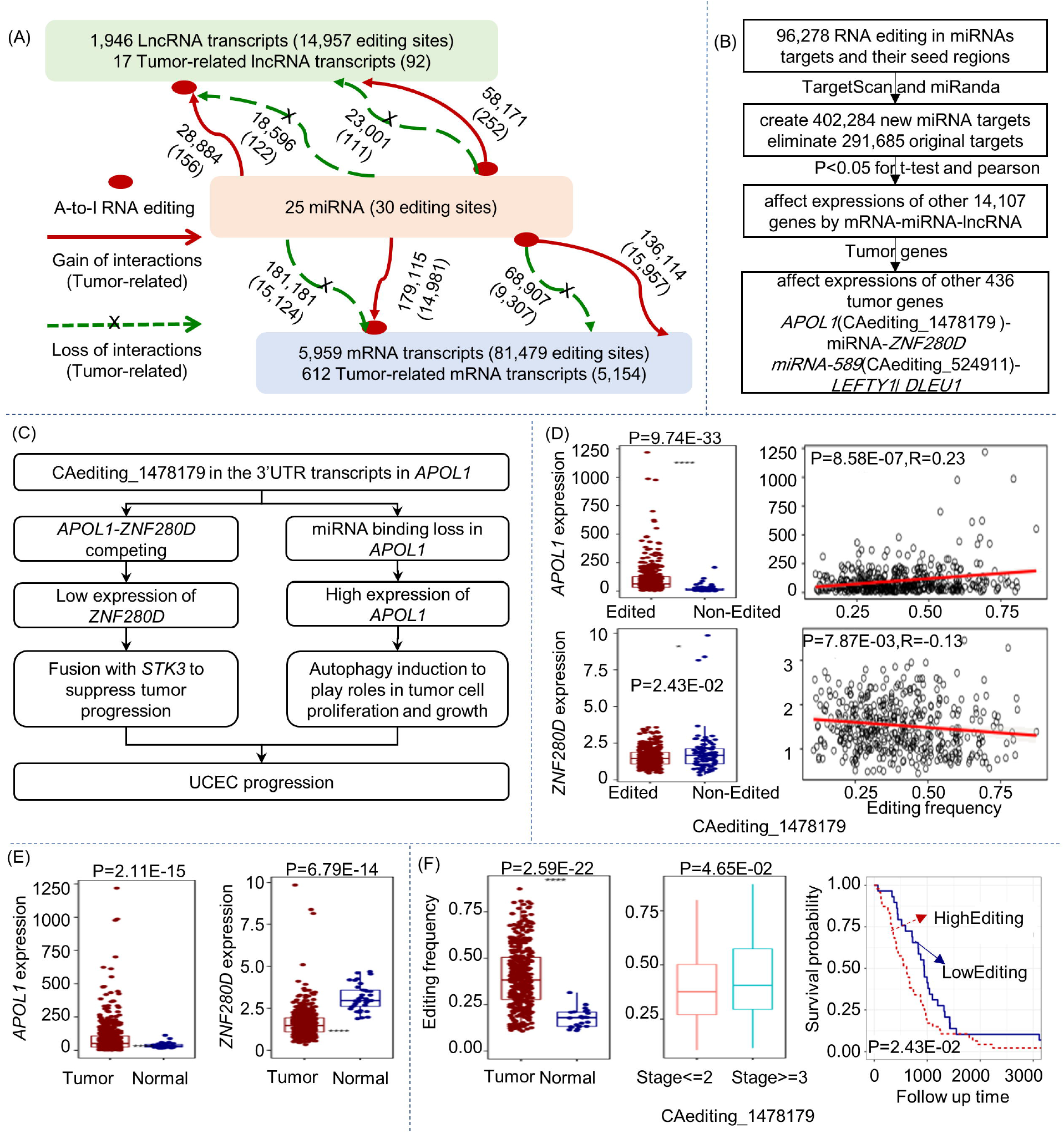
The effects of A-to-I RNA editing events on miRNA regulations. (A) RNA editing events in the 3’-UTRs of mRNAs, lncRNAs, and miRNA seed regions led to the changes of miRNA-targets interactions. (B) The analysis procedures for the effects of A-to-I RNA editing events on miRNA regulations. (C) One significant RNA editing event in the 3’-UTR of *APOL1* likely intervened in the miRNA regulations on two tumor genes to play its roles in the progression of uterine corpus endometrial carcinoma (UCEC). (D) This RNA editing event caused the loss of original miRNA binding target on *APOL1*. The altered miRNA regulation resulted in the increased expressions of *APOL1* and indirectly led to the reduced expressions of competing *ZNF280D* gene. (E) These two genes also showed differential expressions in UCEC, revealing their potential roles and functions in this cancer type. (F) All the analyses uncovered this RNA editing event as a probably pathological biomarker of UCEC, which was also supported by its abnormal editing frequencies in this cancer type. Another significant case shown in panel B was described in Fig. S14.

For example, an RNA editing site (CAediting_1478179) in the 3’-UTR of *APOL1*-201/ 202/ 205/ 206 isoforms would lead to the loss of original binding targets of *miR-7151-3p* (Fig. 6C). The lost regulation seems to cause the increased expressions of *APOL1* and indirectly lead to the reduced expressions of *ZNF280D* with lncRNA transcripts in uterine corpus endometrial carcinoma (UCEC), due to their competing relationships (Fig. 6D). For these two genes, several recent studies reported the induction function of *APOL1* in autophagy (16) to probably promote tumor cell growth and proliferation (36), and the fusion possibility of *ZNF280D* with tumor suppressor gene of *STK3* (37, 38) to involve in cancer. Besides, the DEG analysis also revealed the up-regulation of *APOL1* and down-regulation of *ZNF280D* in the UCEC tumor samples (Fig. 6E). Therefore, we may infer CAediting_1478179 as a progression biomarker for the UCEC cancer type. It was also supported by the significantly higher frequencies of this editing event in the tumor samples, along with more severe tumor statuses, and for the poor survival group (Fig. 6F).

Another RNA editing example located in chr7: 5495852 (CAediting_524911) of *miR-589-3p*. It altered the miRNA binding target from original *DLEU1* to *LEFTY1* (Fig. S14). The lost miRNA regulations led to the over-expressions of *DLEU1*, while the gained interactions caused the down-expressions of *LEFTY1* in the RNA-edited testicular germ cell tumors (TGCT). From previous studies, we found that *DLEU1* is one lncRNA produced from the 13q14.3 tumor suppressor locus and regulates the NF-kB signaling pathway which plays crucial roles in cancer initiation and progression (39–41). We infer a tumor suppressor role of *DLEU1* in TGCT because of its location, functions, and also down-regulations in TGCT tumor samples and that with higher tumor stages. In addition, *LEFTY1*, a key gene in the Nodal pathway, was reported to be specifically associated with germ cell pluripotency, the presence of carcinoma in situ, and TGCT (42). Moreover, this gene was also verified to be up-regulated in TCGT and along with more severe tumor samples. Due to the functions and expression alterations of these two genes, we could speculate that the edited *miR-589-3p* might potentially alleviate TGCT tumor condition, through the up-regulation of *DLEU1* and down-regulation of *LEFTY1*. In conclusion, these two editing candidates represent the potential functions of A-to-I RNA editing events in cancer through altering the miRNA regulations on tumor genes.

## Discussion

Through measuring and analyzing RNA editing events in human pan-cancers, we could provide 24,236 potentially functional A-to-I RNA editing candidates (Table S3). They either were abnormally edited in cancer or might alter the original expression profiles, protein functions, splicing patterns, or miRNA regulations of tumor genes. Considering the contributions of these tumor genes to glutamine metabolism (43, 44), modified immunity (45, 46), selective autophagy (47), DNA damage responses (48), and so on, we may infer that the functional A-to-I RNA editing events may play important roles in tumorigenesis. In the future, the appearance of more functional RNA editing candidates will be accompanied by the increase of tumor genes. The possible RNA editing events and their functions were all archived in CAeditome database.

Among them, five events were explored in detail to introduce their potential functions related to cancer in this study. They are CAediting_390714 of *GluA2*, CAediting_1426931 of *BLCAP*, CAediting_1478179 of *APOL1*, CAediting_543208 of *IGFBP3*, and CAediting_524911 of *miR-589-3p*. The R/G editing in *GluA2* (CAediting_390714) was studied deeply in previous work for its involvements in the desensitization of AMPA receptor (AMPAR) channels (31), AMPAR-mediated neurotransmission, and neurodevelopmental deficits (49–51). For its potentials in cancers, one previous study validated its roles in tumor survival, cell viability, and targeted therapeutics (10). In our study, for this editing event, we discovered anomalously lower editing frequencies in GBM and positive associations with aberrant *GluA2* expression profiles and alternative splicing values in PCPG. Due to the roles of this gene in proliferation stimulation, apoptosis resistance, migration, and invasion in cancer cell lines (19), we may suggest the possible bi-functions of this RNA editing event in neurological and brain tumors. As for the Q/R editing in *BLCAP* (CAediting_1426931), it was abnormally edited in multiple cancer types, such as the mere occurrence in the tumor samples of BLCA, COAD, HNSC, CHOL, and READ, higher editing frequencies in BRCA and KIRC, and positive associations with BLCA tumor stages. The analyses in our study expanded its roles of carcinogenesis promotion in pan-cancers from the cervical cancer reported in previous literature (28). The third RNA editing event (CAediting_1478179) seems to be a novel and promising pathological biomarker for various cancer types including cervical squamous cell carcinoma and endocervical adenocarcinoma (CESC), COAD, esophageal carcinoma (ESCA), and so on. Especially, it was remarkably up-edited in tumors, more severe tumor samples, and poorer survival groups for the cancer types of KIRC (tumor vs. normal: P = 8.08E-05; editing vs. stages: P = 1.71E-10 and R = 0.28; editing vs. survival: P_KM_ = 3.21E-02, P_COX_ = 4.75E-02, and Hazard ratio (HR) = 3.42), lung adenocarcinoma (LUAD, tumor vs. normal: P = 1.52E-25; editing vs. stages: P = 6.59E-03 and R = 0.12; editing vs. survival: P_KM_ = 2.36E-02, P_COX_ = 6.81E-03, and HR = 3.88), and UCEC (tumor vs. normal: P = 2.59E-22; editing vs. stages: P = 4.32E-02 and R = 0.10; editing vs. survival: P_KM_ = 2.43E-02, P_COX_ = 1.50E-02, and HR = 5.74). Moreover, we discovered that it caused the loss of original miRNA binding targets to potentially induce the over-expressions of *APOL1*, which was supported by their positive associations in pan-cancers, such as for KIRC (P = 7.44E-06 and R = 0.20), LUAD (P = 6.78E-10 and R = 0.27), and UCEC (P = 8.58E-07 and R = 0.23). Thus, this RNA editing event may confer its pathological function in cancers through its intervention in miRNA regulations on the tumor gene of *APOL1*. Another novel RNA editing event (CAediting_543208) occurred only in the tumor samples for the KIRC cancer type. It may enhance the inhibition of tumor cell growth through its positive impacts on the tumor suppressor gene of *IGFBP3* (17, 18). The last RNA editing event located (CAediting_524911) in the seed region of *miR-589-3p* to potentially modify its original regulations on many tumor genes. For example, the edited *miR-589-3p* altered the expression levels of *DLEU1* and *LEFTY1*, which thus may alleviate TGCT tumor condition. Moreover, another evidence in one previous study also validated the regulatory potential of this RNA editing event (CAediting_524911) on two genes of *PCDH9* and *ADAM12* to control glioblastoma cell migration and invasion (52).

Of the five edited genes introduced in this study, three were discovered to be linked with tumor-related phenotypes from DisGeNet (January 2021, version 7.0) (53). Specifically, *APOL1* is associated with neoplasm-related nephrotic syndrome (45) and common focal segmental glomerulosclerosis form of kidney disease (46), *IGFBP3* plays important roles in multiple cancers (54), and *GluA2* is related to neurological or brain diseases (53). Their possible relationships with these disorders may be partially attributed to the RNA editing events in them, which may also affect the effectiveness of probable drugs targeting them. In total, we discovered 6717 edited genes associated with 9203 different types of diseases in DisGeNet, and 1586 edited genes targeted by 1674 approved drugs from DrugBank (January 2021, version 5.1.8) (55). The functions of A-to-I RNA editing events in these genes will be useful for exploring the pathological mechanisms of related diseases and providing novel knowledge to design the targeted drugs.

Beside the five RNA editing events introduced in detail, there are also several potential candidates archived in CAeditome database which were validated or proposed in previous studies. For example, in our study, the analysis of RNA editing events in protein-coding regions identified I164V in *COPA* (CAediting_115738), S367G in *AZIN1* (CAediting_655260), and I635V in *COG3* (CAediting_962851). These three RNA editing events were also abnormally edited in tumor samples, correlated with tumor severity, associated with cancer survival, and possible factors to affect the expressions of their host genes in multiple cancer types. Their functions in tumorigenesis have been validated in cell lines and mice models by other groups (10, 56–61). In addition, the analysis of RNA editing events in miRNA seed regions uncovered altered *miR-200b* regulations associated with CAediting_393. This event leads to the gain/ loss of many miRNA binding targets, including that in *LIFR, ZEB1*, and *ZEB2*. Although the expressions of these three genes were not significantly associated with the frequencies of this editing event as shown in different cell lines previously (8, 62), we still included it as a potentially functional A-to-I RNA editing candidate, since it probably dys-regulated other tumor-related genes (Download Page in CAeditome database). For example, *ACSL6* was down-regulated to possibly interfere in the metabolites of fatty acids, abnormality of which is one of the cancer hallmarks (63). Another gene of *PKP1* was inhibited to prevent the survival and metastasis of cancer cells by decreasing cluster formation in circulatory system (64). Moreover, there are also other RNA editing events whose functions in cancers were bio-experimentally validated, including H241R editing (CAediting_604339) in *PODXL* (5), K242R/K242E (CAediting_1062027, CAediting_1062028) editing in *NEIL1* (65), two editing events (CAediting_442015, CAediting_442019) in the 3’-UTR of *GM2A* (66), and so on. The other potential RNA editing biomarkers are waiting for the validation by cancer research communities.

In summary, this study proposed a transcriptome-wide and cancer-wide map for the functions of individual A-to-I RNA editing event. It will provide the chances to understand cancer pathology from the A-to-I RNA editing aspect and list potential biomarkers and therapeutic targets for cancer and drug research communities. However, during the analyses, the complex regulatory mechanisms associated with RNA editing pointed out two possible studies in the future.

Firstly, we noted the possible interactions of multiple RNA editing events and their co-regulatory effects on the downstream genes or regulations. For example, three RNA editing events were all in together to confer their effects on the expressions of *APOL1*, which were deciphered by the least absolute shrinkage and selection operator regression method as shown in Fig. S15. In the future, to uncover the co-regulations of RNA editing events, we will propose an RNA editing weighted gene expression network and evaluate its usefulness in various clinical scenarios such as survival prognosis.

On the other hand, we confirmed the deamination functions of three editing enzymes on 11,948 RNA editing events in this study (P < 0.05 and R > 0.3, Fig. S16). However, we should not ignore that there are another 28,062 RNA editing events showing no statistical associations with all the three enzymes. It revealed the possible multi-regulators of A-to-I RNA editing. From previous literature, these diverse regulatory mechanisms probably include genetic variations (67, 68), splicing efficiency (69, 70), and RNA binding proteins (71, 72).

For a functional RNA editing candidate (CAediting_1478179 of *APOL1)* proposed in this study, we also discovered its differential editing frequencies among the genotyping groups of three single nucleotide variations in the KIRC cancer type (Fig. S17). For the impacts of genetic variations on RNA editing events, recently, Leng Han group has published a database named GPEdit (73). Then the studies on the other potential regulators of A-to-I RNA editing are our further research plans.

## Materials and methods

### Detection of A-to-I RNA editing

For all the 11,056 samples across 33 cancer types in TCGA (Supplementary Table 1), we first detected RNA editing events from individual RNA-seq bam files by the script of REDItoolKnown.py (REDItools 1.2.1) (74) with default settings (e.g., minimal read coverage, 10; minimal quality score, 30; and minimal mapping quality score, 255) and hg38 (GENCODE (v.22) reference files same as that used in the GDC data harmonization and generation pipelines. To ensure the confident identification of RNA editing events, we focused on known editing sites from REDIportal (January 2021) (75), removed possible SNP data (dbSNP151 and genome-wide human SNP array 6.0), and filtered out the candidates with supporting reads under three or editing frequencies less than 0.1. Eventually, we selected one kind of RNA editing types, A-to-I RNA editing for further analysis, because of its abundance in humans.

For all the detected A-to-I RNA editing events, we analyzed their distributions in diverse genomic locations and repeats by ANNOVAR (76) and evaluated the stability alterations of edited transcripts by RNAfold (ViennaRNA 2.4.17) (77). The consistency of these analysis results (Fig. S18-S19) with previous publications (13, 78, 79) revealed the reliability of A-to-I RNA editing detection in this study. Moreover, all these events covered a high ratio of RNA editing sites (72.49%) detected in one previous study (10) as shown in Fig. S20. It describes the consistency of our RNA editing detection pipeline with others. In addition, the more RNA editing sites detected in our study provided an opportunity for a more comprehensive map of functional A-to-I RNA editing events.

### Analysis of A-to-I RNA editing frequencies

To uncover potential A-to-I RNA editing events related to tumors, we first compared their editing frequencies between tumor samples and controls across 33 cancer types. Then we defined a tumor-specific RNA editing event if it only occurred in tumors with more than 5 edited samples or showed significantly differential editing frequencies (P < 0.05) in tumor samples. Next, we analyzed the correlations between editing frequencies and tumor stages (pathologic stage or clinical stage, P < 0.05), to identify tumor progression-associated RNA editing events. Third, we performed Kaplan–Meier (KM), and Cox Proportional-Hazards regression (COX) analyses to determine the A-to-I RNA editing events (P < 0.05 for both results) which may affect tumor survival. Last, we focused on the RNA editing events in 1615 tumor-related genes, including driver oncogenes from OncoVar (80), tumor suppressor genes from TSGene 2.0 (81), and some other genes reported to be associated with tumors in previous literature (16, 82), in order to further analyze the possible effects of A-to-I RNA editing in cancer.

### Analyzing A-to-I RNA editing effects on gene expressions and pathways

To study the effects of A-to-I RNA editing events on gene expressions, we performed the analyses of differentially expressed genes (DEGs) between RNA-edited and non-edited tumor samples (P < 0.05 and |log2FC| > 0.3) by T test and Pearson correlations between editing frequencies and corresponding gene expressions (P < 0.05) across 33 cancer types. The dys-regulated genes in RNA-edited groups and also along with the changes of editing frequencies were inferred to be potentially affected by A-to-I RNA editing. These genes combined with the DEGs in tumors compared to controls (T test: P < 0.05 and |log2FC| > 0.3) were further studied by Enrichr (83), to assess the probably involved cellular processes of A-to-I RNA editing in cancer.

### Analyzing A-to-I RNA editing effects on protein recoding and functions

For A-to-I RNA editing events in coding regions, we used ANNOVAR to detect the changes of amino acid sequences caused by the non-synonymous and stop-loss editing sites. Then their deleterious effects on protein functions were assessed by SIFT, Polyphen2, and PROVEAN (dbNSFP version 4.1a).

### Analyzing A-to-I RNA editing effects on alternative splicing of pre-mRNAs

To study the effects of A-to-I RNA editing events on splicing, we first overlapped them with 5’-donor splice sites (5’-ss) and 3’-acceptor splice sites (3’-ss) around detected exons (84). The 5’-ss is a 9-mer region of 3 nucleotides (nt) in the exon and 6 nt in the intron, while 3’-ss is a 23-mer region of 3 nt in the exon and 20 nt in the intron based on a previous splicing study (85). MaxEntScan method proposed in that study was also used here to estimate the changes of splice site strength for sequences being edited. These splicing alterations were further validated by the comparisons of PSI values between RNA-edited and non-edited tumor samples (P < 0.05) and the correlations of PSI values with corresponding editing frequencies (P < 0.05), to discover the reliable effects of A-to-I RNA editing events on splicing patterns.

### Analyzing A-to-I RNA editing effects on miRNA regulations

For the wild-type and RNA-edited 3’-UTRs of mRNA, lncRNAs, and miRNA seed regions, we used TargetScan (14) (version: 7.0) and miRanda (15) (version: 3.3a) to detect miRNA binding targets. Based on the predicted miRNA–lncRNA/ mRNA 3’-UTR interactions, we defined the gain of miRNA binding targets as the interactions existing in the RNA-edited sequences but not in the wild-type sequences supported by both tools and vice versa for the loss of miRNA binding targets. Furthermore, we checked the expressions of miRNA-regulated genes between RNA-edited and non-edited tumor groups (P < 0.05) and also along with the changes of editing frequencies (P < 0.05), to discover the altered miRNA regulations caused by these A-to-I RNA editing events.

## Credit author statement

**Sijia Wu:** Conceptualization, Data Curation, Methodology, Investigation, Software, Formal analysis, Writing - Original Draft, Funding acquisition. **Zhiwei Fan:** Visualization. **Pora Kim:** Conceptualization, Visualization, Validation, Writing - Review & Editing. **Liyu Huang:** Supervision, Project administration, Resources, Funding acquisition. **Xiaobo Zhou:** Supervision, Project administration, Resources

## Acknowledgements

The results here are in whole or part based upon data generated by the TCGA Research Network in https://www.cancer.gov/tcga. This work was supported by the National Natural Science Foundation of China (No. 62002270), the Fundamental Research Funds for the Central Universities, the Natural Science Foundation of Shaanxi Province of China (No. 2020JQ-332), the China Postdoctoral Science Foundation (No. 2018M643583), National Key Research and Development Program of China (Grant No. 2017YFA0205202), and partially funded by the National Natural Science Foundation of China (Grant No.61672422). The funders had no role in study design, data collection and analysis, decision to publish or preparation of the manuscript.

## Supplementary material

**Supplementary Figure 1 Tumor-specific RNA editing events with more than 5 edited tumor samples and none edited normal controls.**

The x-axis indicates the number of edited tumor samples, and the y-axis represents the number of editing sites. The genes shown in this figure are tumor-related genes with this kind of tumor-specific RNA editing events. Specifically, CAediting_543208 (chr7:45916046) of *IGFBP3* occurred only in tumor samples (246/535) and none in controls (0/72) for kidney renal clear cell carcinoma. The 33 cancer types involved in this study included adrenocortical carcinoma (ACC), bladder urothelial carcinoma (BLCA), breast invasive carcinoma (BRCA), cervical squamous cell carcinoma and endocervical adenocarcinoma (CESC), cholangiocarcinoma (CHOL), colon adenocarcinoma (COAD), lymphoid neoplasm diffuse large b-cell lymphoma (DLBC), esophageal carcinoma (ESCA), glioblastoma multiforme (GBM), head and neck squamous cell carcinoma (HNSC), kidney chromophobe (KICH), kidney renal clear cell carcinoma (KIRC), kidney renal papillary cell carcinoma (KIRP), acute myeloid leukemia (LAML), brain lower grade glioma (LGG), liver hepatocellular carcinoma (LIHC), lung adenocarcinoma (LUAD), lung squamous cell carcinoma (LUSC), mesothelioma (MESO), ovarian serous cystadenocarcinoma (OV), pancreatic adenocarcinoma (PAAD), pheochromocytoma and paraganglioma (PCPG), prostate adenocarcinoma (PRAD), rectum adenocarcinoma (READ), sarcoma (SARC), skin cutaneous melanoma (SKCM), stomach adenocarcinoma (STAD), testicular germ cell tumors (TGCT), thyroid carcinoma (THCA), thymoma (THYM), uterine corpus endometrial carcinoma (UCEC), uterine carcinosarcoma (UCS), and uveal melanoma (UVM).

**Supplementary Figure 2 The bubble plots of top tumor-specific RNA editing events (edited only in tumor samples with number ≧ 5) in tumor genes for each cancer type.**

The x-axis shows the number of edited tumor samples, and the y-axis represents the editing events and their host tumor genes

**Supplementary Figure 3 The volcano plots for tumor-specific RNA editing events (P < 0.05 and edited samples ≧ 50).**

The x-axis denotes the differences of editing frequencies between tumor samples and controls, and the y-axis represents −log10(P) value. The genes shown in this figure are tumor-related genes with this kind of tumor-specific RNA editing events.

**Supplementary Figure 4 The bubble plots of top tumor-specific RNA editing events (P < 0.05 and edited samples ≧ 50) in tumor genes for each cancer type.**

The x-axis shows log10(P) value, and the y-axis represents the editing events and their host tumor genes. The size of each dot denotes the absolute differences of editing frequencies. The red and green dots represent the RNA editing events showing significantly higher or lower frequencies in tumor samples.

**Supplementary Figure 5 The volcano plots for tumor stage-associated RNA editing events (P < 0.05 and edited tumor samples ≧ 50).**

The x-axis denotes the correlation coefficients between editing frequencies and tumor stages, and the y-axis represents −log10(P) value. The genes shown in this figure are tumor-related genes with tumor stage-associated RNA editing events.

**Supplementary Figure 6 The bubble plots of top tumor stage-associated RNA editing events (P < 0.05 and edited tumor samples ≧ 50) in tumor genes for each cancer type.**

The x-axis shows log10(P) value, and the y-axis represents the editing events and their host tumor genes. The size of each dot denotes the absolute correlation coefficients. The red and green dots represent the RNA editing events showing positive or negative relationships with tumor stages.

**Supplementary Figure 7 The volcano plots for tumor survival-related RNA editing events (P < 0.05 and edited tumor samples ≧ 50).**

If Hazard ratio (HR) is smaller than 1.0, then the x-axis denotes the value of HR-1. Otherwise, the x-axis denotes the value of (HR-1)/HR. The y-axis represents −log10(P) value. The genes shown in this figure are tumor-related genes with tumor survival-related RNA editing events.

**Supplementary Figure 8 The bubble plots of top tumor survival-related RNA editing events (P < 0.05 and edited tumor samples ≧ 50) in tumor genes for each cancer type.**

The x-axis shows log10(P) value, and the y-axis represents the editing events and their host tumor genes. The size of each dot denotes −log10(P) value. The red and green dots represent the RNA editing events showing high (HR > 1) or low (HR < 1) survival risks for cancer patients.

**Supplementary Figure 9 The potential of CAediting_543208 *(IGFBP3)* in KIRC cancer type.**

This event occurred only in tumor samples (246/535) and none in controls (0/72), and was up-edited in tumor samples with higher stages. It was also related to the over-expressions of *IGFBP3*, which was up-regulated in tumor samples and appeared to act in an autocrine action to suppress tumor cell growth. Thus, this RNA editing event might enhance the protection functions of *IGFBP3* against cancer progression and be potential as a therapeutic target.

**Supplementary Figure 10 The volcano plots for DEG-associated RNA editing events in each cancer type.**

These events were identified according to the procedures shown in Fig. 3E. The x-axis denotes log2(Fold change) value between RNA-edited and non-edited tumor samples, and the y-axis represents −log10(P) value. The genes shown in this figure are tumor-related genes with DEG-associated RNA editing events.

**Supplementary Figure 11 The bubble plots of top DEG-associated RNA editing events in tumor genes for each cancer type.**

The x-axis shows log10(P) value, and the y-axis represents the editing events and their host tumor genes. The size of each dot denotes absolute log2(Fold change) value between RNA-edited and non-edited tumor samples. The red and green dots represent the RNA editing events positively or negatively associated with the expressions of their host genes.

**Supplementary Figure 12 An example (chr4:157360142, CAediting_390714) showing the effects of RNA editing on gene expressions.**

The frequencies of this editing event were positively associated with the expressions of its host gene *(GRIA2)* in PCPG. Since this editing event led to the significantly higher expressions of its host gene, which was also up regulated in tumor samples, CAediting_390714 may be a pathological biomarker for the PCPG cancer type. It was also supported by 146 (183) edited tumor samples and 0 (3) edited controls in this cancer type.

**Supplementary Figure 13 The enriched KEGG pathways of the overlapped DEGs for each cancer type (p < 0.05 and q < 0.2).**

The pathways colored with brown are tumor-related pathways.

**Supplementary Figure 14 An example showing the effects of RNA editing on miRNA regulations.**

An RNA editing event in chr7:5495852 (CAediting_524911) of *miR589-3p* altered the miRNA binding target from original *DLEU1* to *LEFTY1*. Due to the lost regulation of *miR-589-3p, DLEU1* was up-regulated in the RNA-edited TGCT tumor samples. On the other hand, *LEFTY1* was down-regulated because of the gained interactions with *miR-589-3p*.

**Supplementary Figure 15 An example showing the co-effects of three RNA editing events on the expressions of *APOL1* in the KIRC cancer type.**

(A) The formula to estimate the co-effects of multiple RNA editing events on gene expressions. (B) The process of least absolute shrinkage and selection operator (Lasso) to calculate the contributions of three RNA editing events to *APOL1* expressions. (C) Contribution coefficients of the three RNA editing events to *APOL1* expressions.

**Supplementary Figure 16 The correlations of RNA editing frequencies with *ADAR* expressions.**

The frequencies of 9702, 2476, and 1919 RNA editing events are significantly associated with the expressions of *ADAR1, ADAR2*, and *ADAR3* respectively (P < 0.05 and R > 0.3 for Pearson method). In addition, there are also 28,062 RNA editing events (P ≧ 0.05 for all three *ADARs* with more than 50 edited samples) which were possibly regulated by genetic variations, splicing efficiency, RNA binding proteins, or some other mechanisms.

**Supplementary Figure 17 The potentially regulatory effects of rs2003814 (A), rs9607326 (B), and rs9607325 (C) on CAediting_1478179.**

**Supplementary Figure 18 The distributions of A-to-I RNA editing events in different types of genes (A), regions (B), and repeats (C).**

**Supplementary Figure 19 The effects of RNA editing on secondary structure.**

(A) An A-to-I RNA editing event will significantly reduce the minimum free energy (MFE) to stabilize RNA structure. (B) 89.32% RNA transcripts were stabilized by one or multiple RNA editing events.

**Supplementary Figure 20 The comparisons of RNA editing events detected by different pipelines.**

For comparison, we converted the genome coordinates of RNA editing events detected in previous study (PMID: 26439496) from GRCh37/hg19 to GRCh38/hg38.

**Supplementary Table 1 The statistical results for RNA editing related cancer mechanisms.**

**Supplementary Table 2 The tumor-related KEGG pathways.**

**Supplementary Table 3 The potentially functional A-to-I RNA editing events.**

