## Supplementary Figure 1-20 for "The Integrative Studies on the Functional A-to-I RNA Editing Events in Human Cancers"

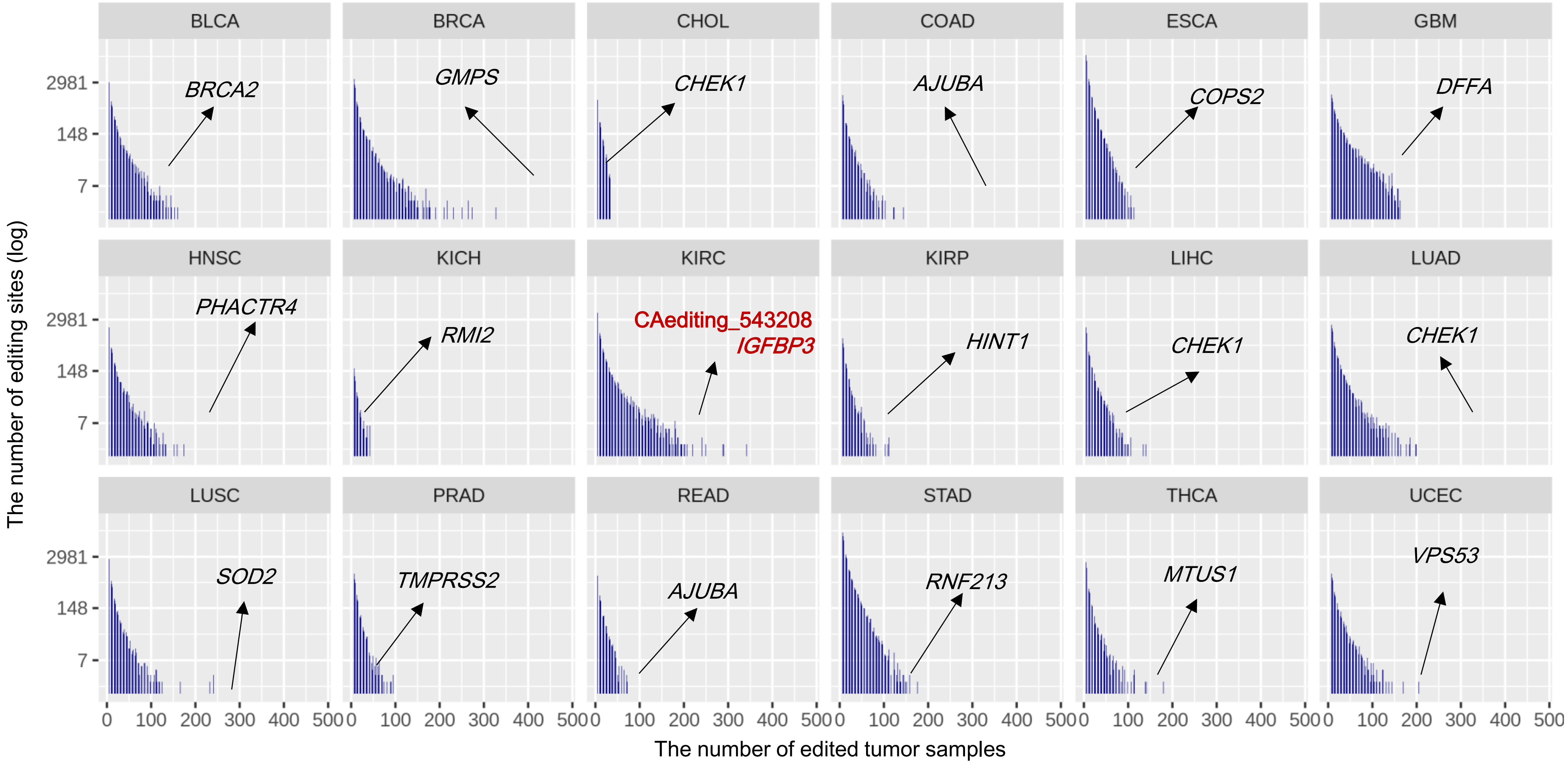


**Fig. S1. Tumor-specific RNA editing events with more than 5 edited tumor samples and none edited normal controls**. The x-axis indicates the number of edited tumor samples, and the y-axis represents the number of editing sites. The genes shown in this figure are tumor-related genes with this kind of tumor-specific RNA editing events. Specifically, CAediting_543208 (chr7:45916046) of *IGFBP3* occurred only in tumor samples (246/535) and none in controls (0/72) for kidney renal clear cell carcinoma. The 33 cancer types involved in this study included adrenocortical carcinoma (ACC), bladder urothelial carcinoma (BLCA), breast invasive carcinoma (BRCA), cervical squamous cell carcinoma and endocervical adenocarcinoma (CESC), cholangiocarcinoma (CHOL), colon adenocarcinoma (COAD), lymphoid neoplasm diffuse large b-cell lymphoma (DLBC), esophageal carcinoma (ESCA), glioblastoma multiforme (GBM), head and neck squamous cell carcinoma (HNSC), kidney chromophobe (KICH), kidney renal clear cell carcinoma (KIRC), kidney renal papillary cell carcinoma (KIRP), acute myeloid leukemia (LAML), brain lower grade glioma (LGG), liver hepatocellular carcinoma (LIHC), lung adenocarcinoma (LUAD), lung squamous cell carcinoma (LUSC), mesothelioma (MESO), ovarian serous cystadenocarcinoma (OV), pancreatic adenocarcinoma (PAAD), pheochromocytoma and paraganglioma (PCPG), prostate adenocarcinoma (PRAD), rectum adenocarcinoma (READ), sarcoma (SARC), skin cutaneous melanoma (SKCM), stomach adenocarcinoma (STAD), testicular germ cell tumors (TGCT) , thyroid carcinoma (THCA), thymoma (THYM), uterine corpus endometrial carcinoma (UCEC), uterine carcinosarcoma (UCS), and uveal melanoma (UVM).

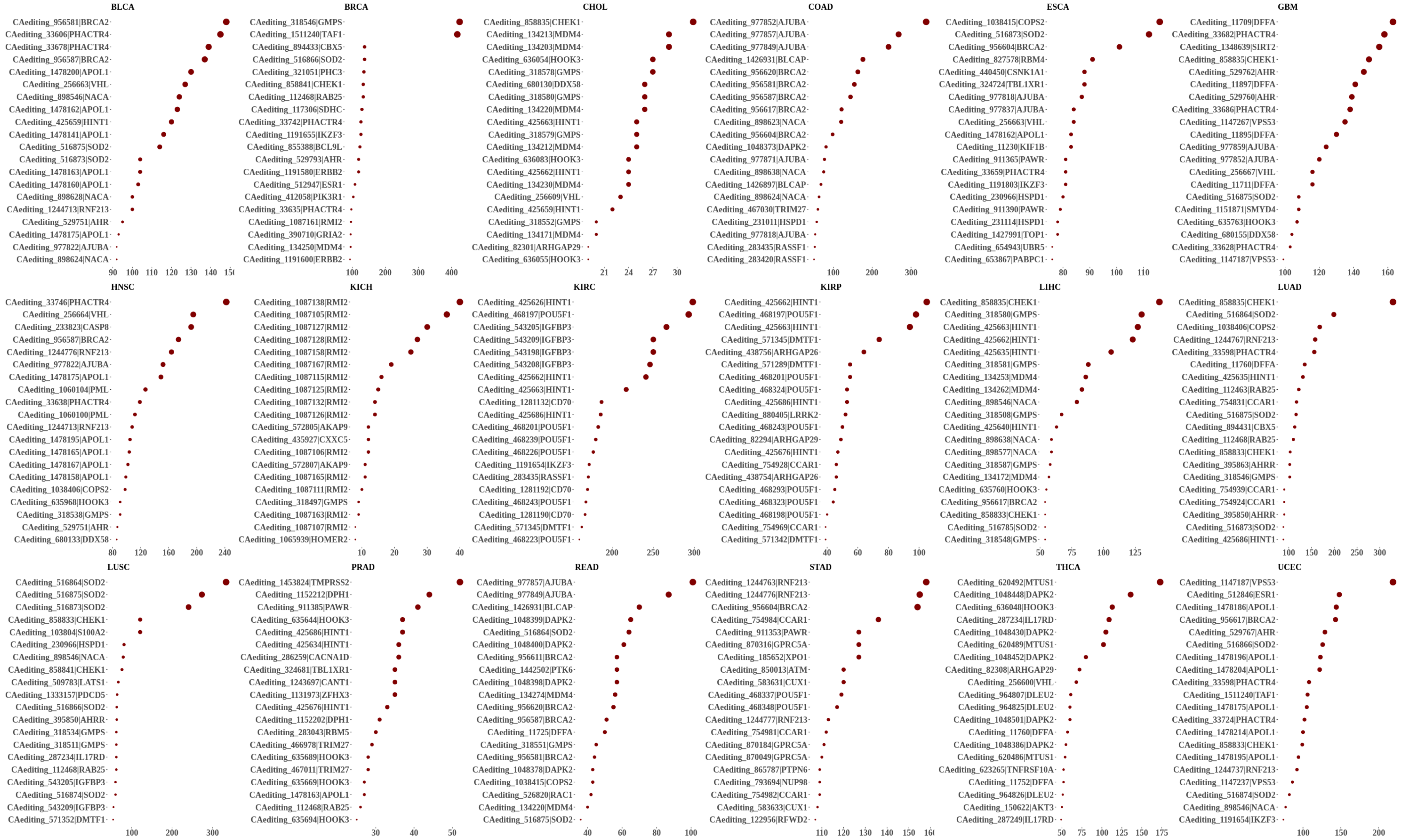


**Fig. S2. The bubble plots of top tumor-specific RNA editing events (edited only in tumor samples with number ≧ 5) in tumor genes for each cancer type.** The x-axis shows the number of edited tumor samples, and the y-axis represents the editing events and their host tumor genes.

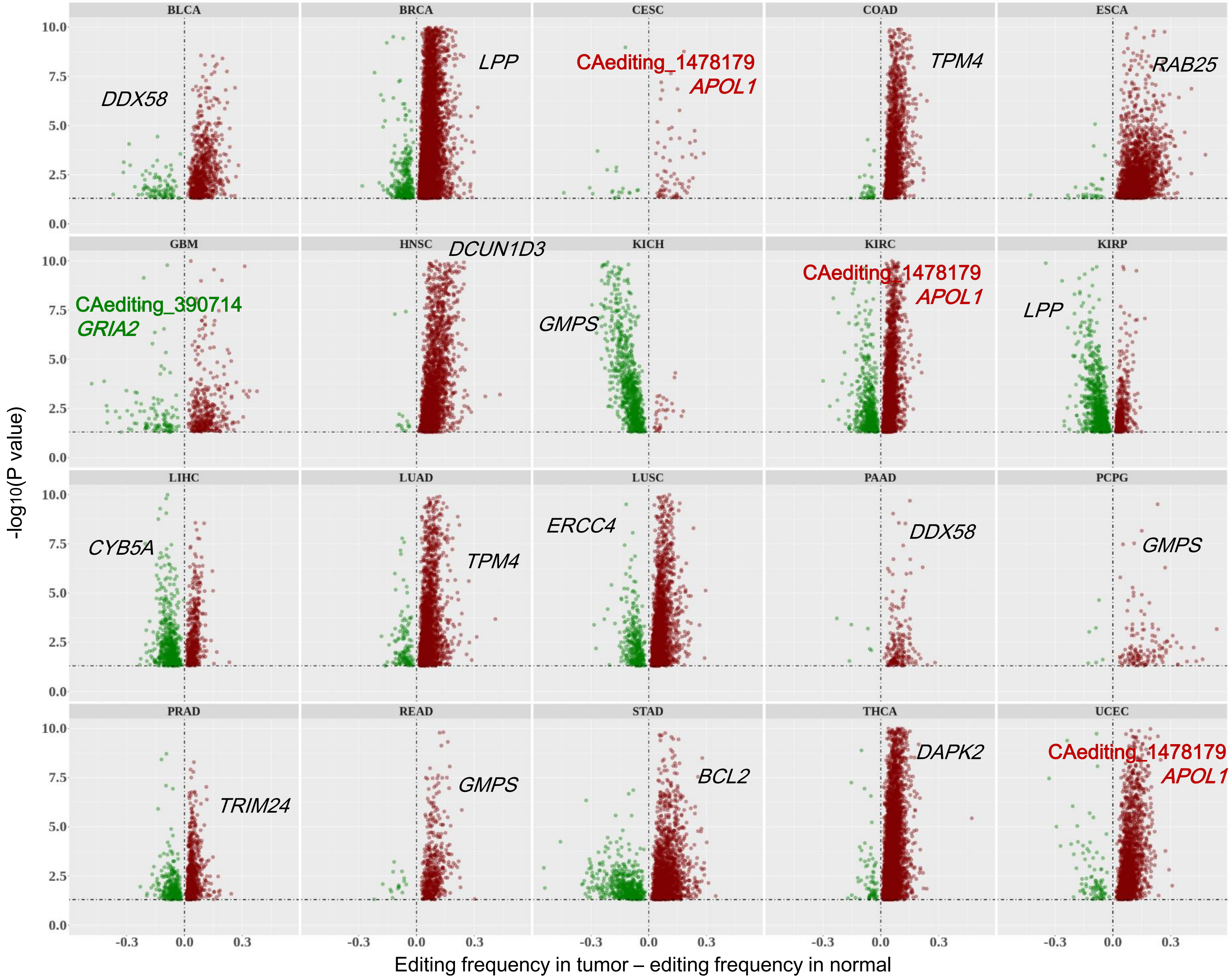


**Fig. S3. The volcano plots for tumor-specific RNA editing events (P < 0.05 and edited samples ≧ 50).** The x-axis denotes the differences of editing frequencies between tumor samples and controls, and the y-axis represents -log10(P) value. The genes shown in this figure are tumor-related genes with this kind of tumor-specific RNA editing events.

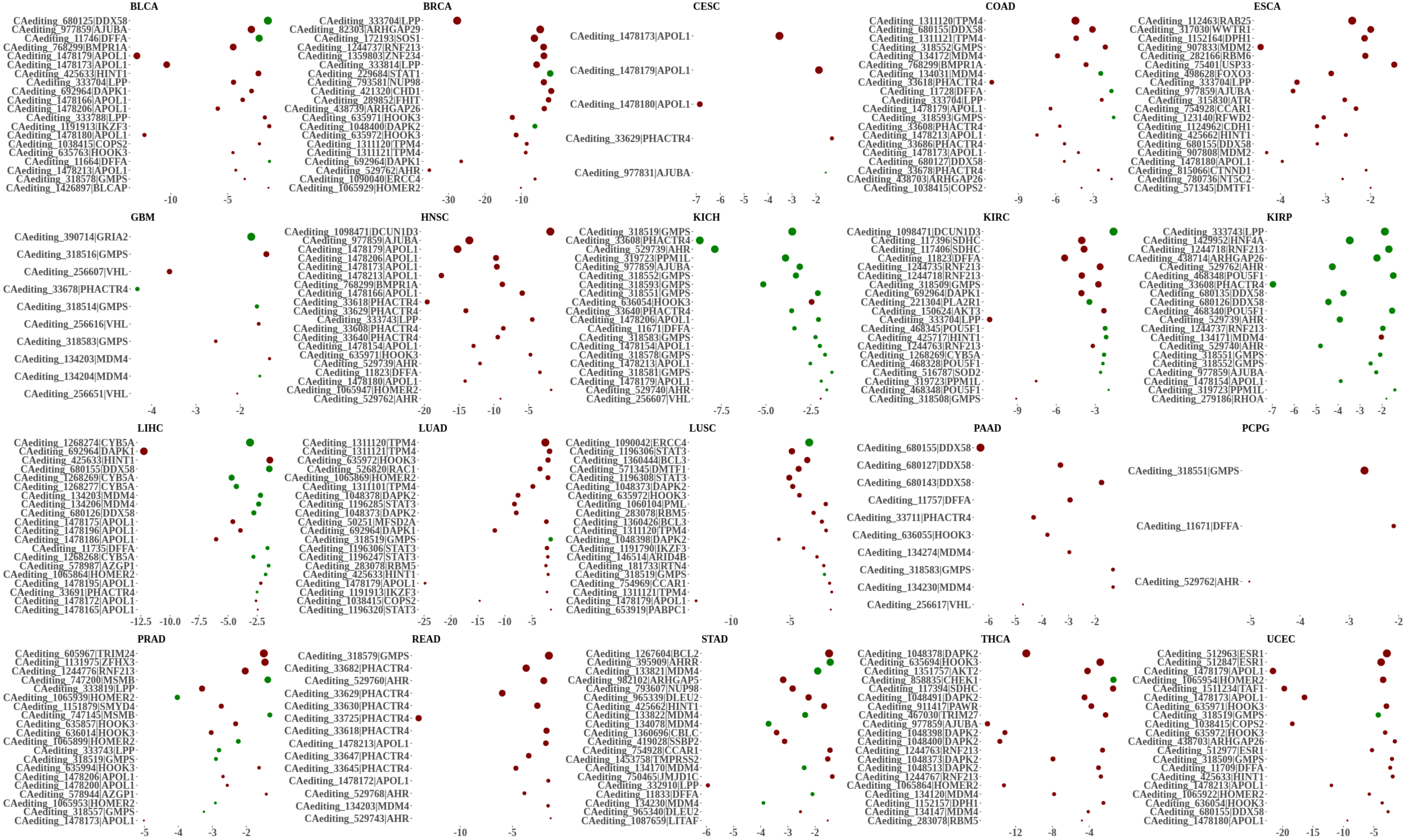


**Fig. S4. The bubble plots of top tumor-specific RNA editing events (P < 0.05 and edited samples ≥ 50) in tumor genes for each cancer type.** The x-axis shows log10(P) value, and the y-axis represents the editing events and their host tumor genes. The size of each dot denotes the absolute differences of editing frequencies. The red and green dots represent the RNA editing events showing significantly higher or lower frequencies in tumor samples.

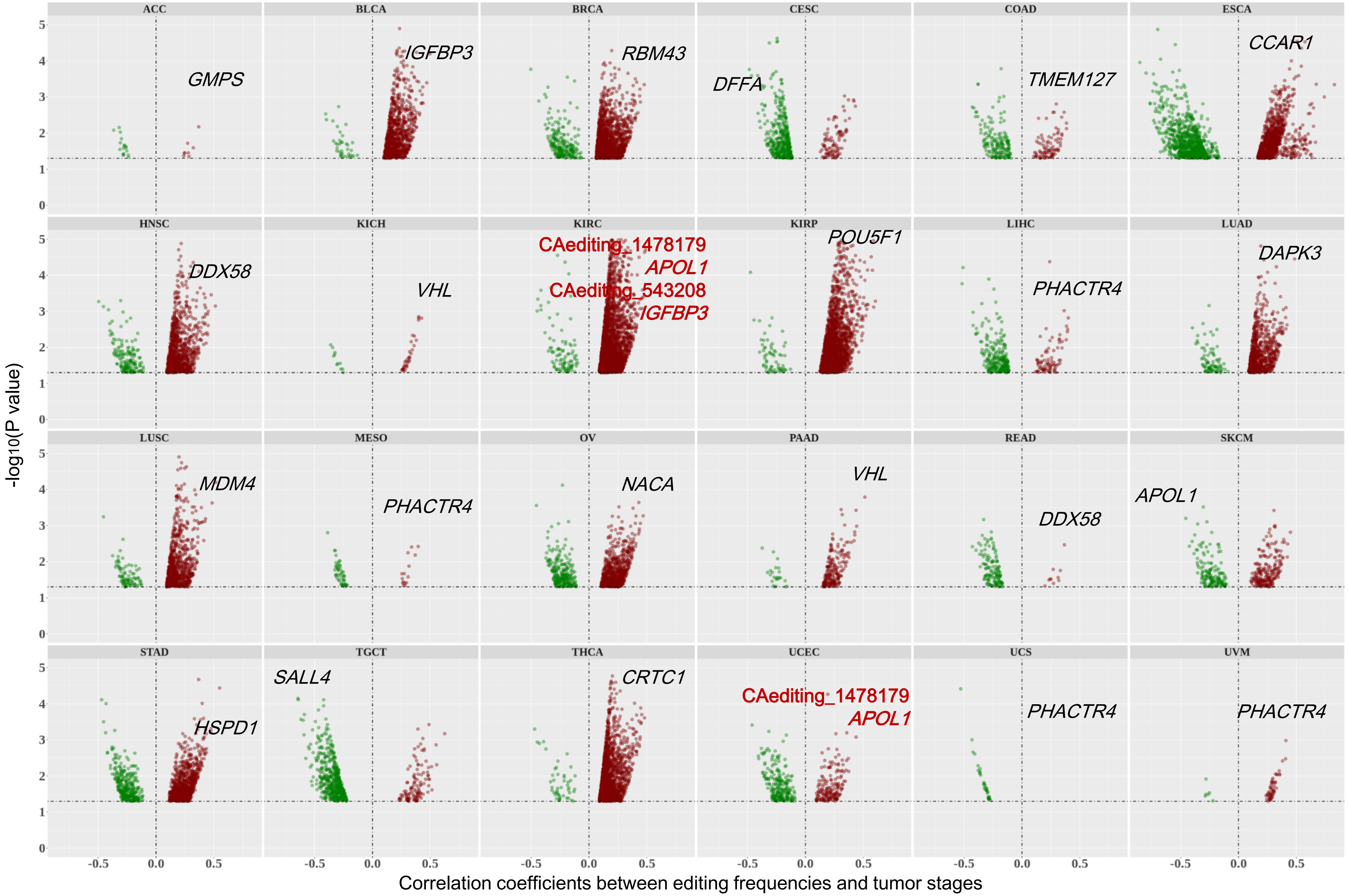


**Fig. S5. The volcano plots for tumor stage-associated RNA editing events ($P < 0.05$ and edited tumor samples ≧ 50).** The x-axis denotes the correlation coefficients between editing frequencies and tumor stages, and the y-axis represents $-\log_{10}(P)$ value. The genes shown in this figure are tumor-related genes with tumor stage-associated RNA editing events.

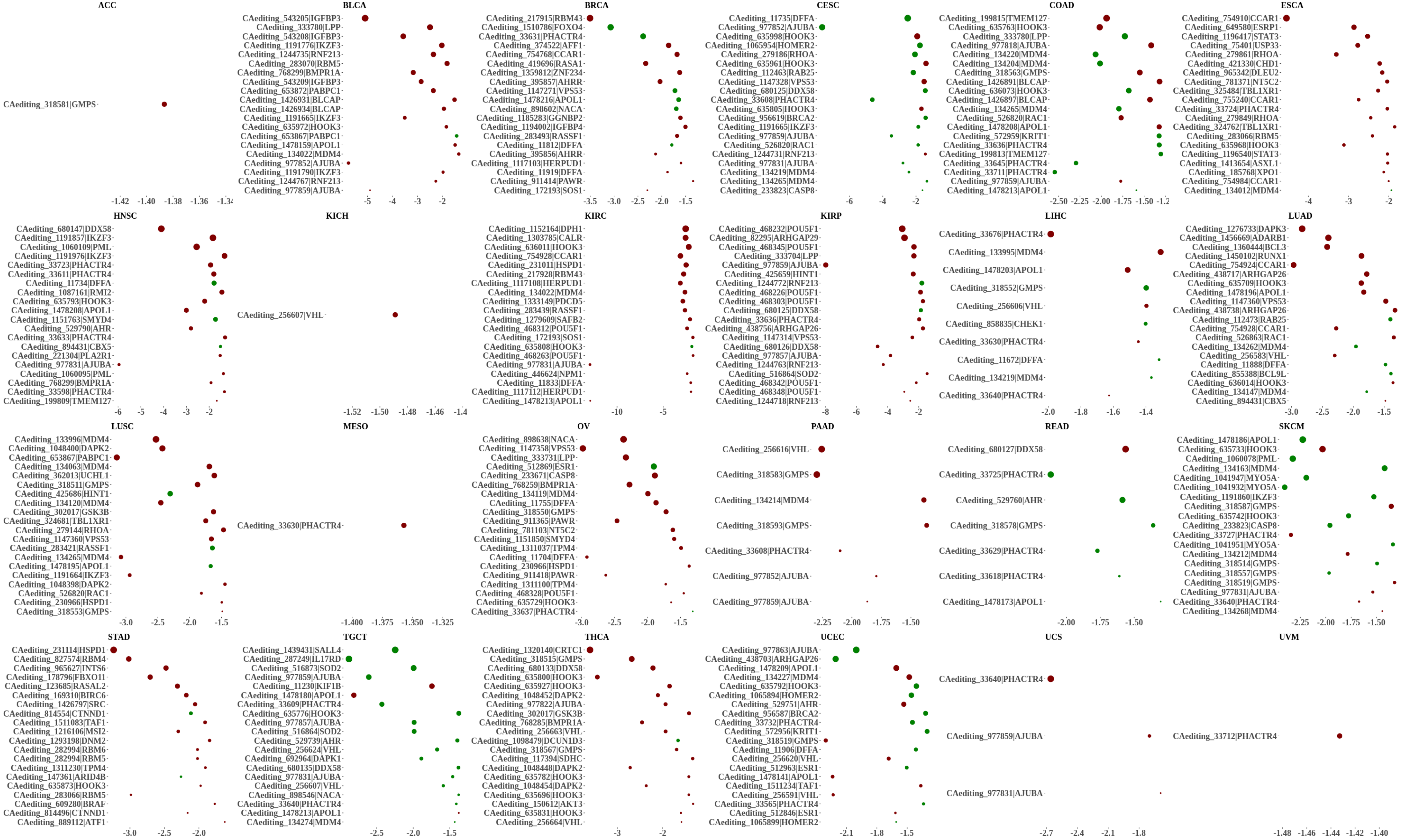


**Fig. S6. The bubble plots of top tumor stage-associated RNA editing events (P < 0.05 and edited tumor samples ≧ 50) in tumor genes for each cancer type.** The x-axis shows log10(P) value, and the y-axis represents the editing events and their host tumor genes. The size of each dot denotes the absolute correlation coefficients. The red and green dots represent the RNA editing events showing positive or negative relationships with tumor stages.

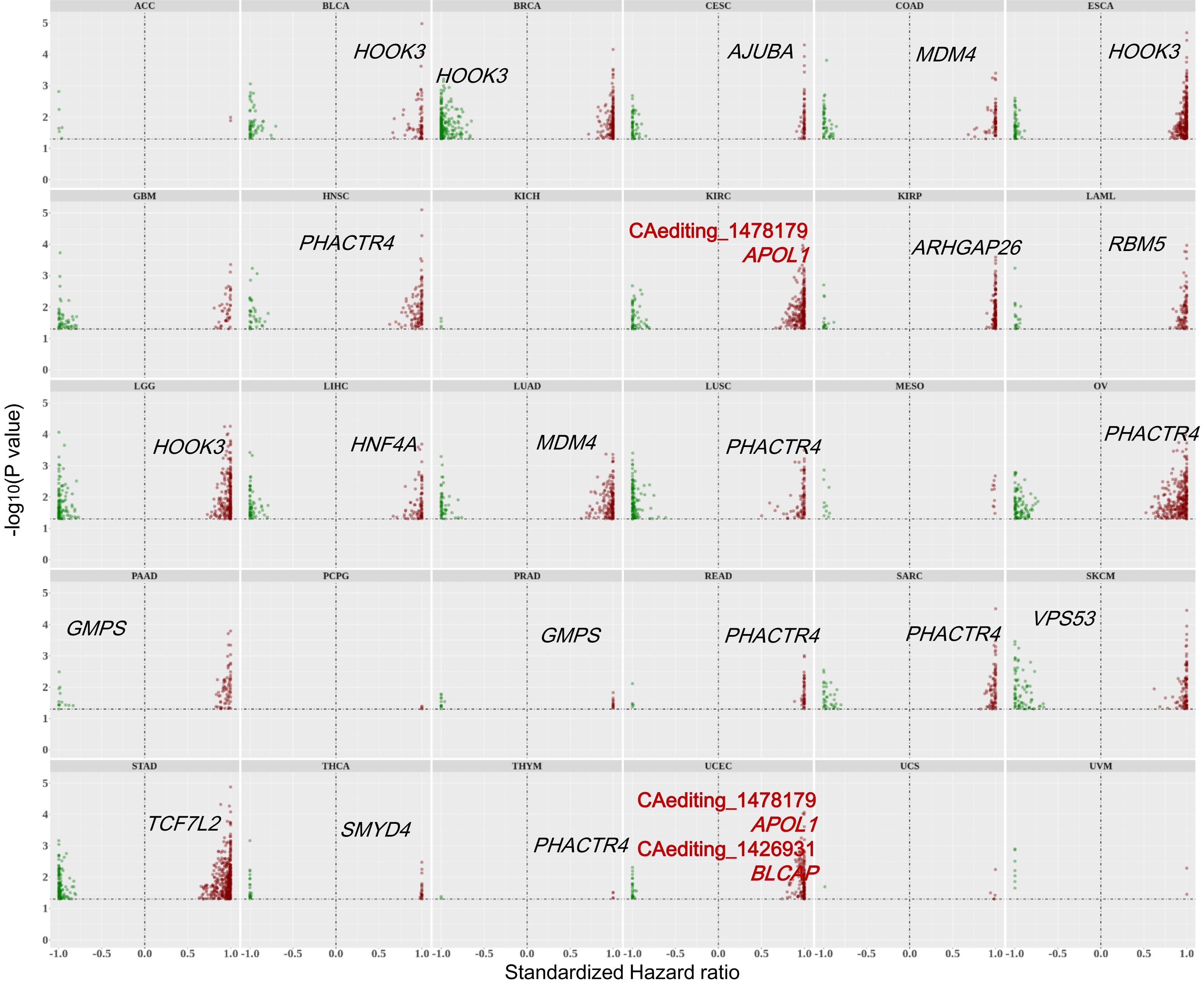


**Fig. S7. The volcano plots for tumor survival-related RNA editing events (P < 0.05 and edited tumor samples ≧ 50).** If Hazard ratio (HR) is smaller than 1.0, then the x-axis denotes the value of HR-1. Otherwise, the x-axis denotes the value of (HR-1)/HR. The y-axis represents -log10(P) value. The genes shown in this figure are tumor-related genes with tumor survival-related RNA editing events.

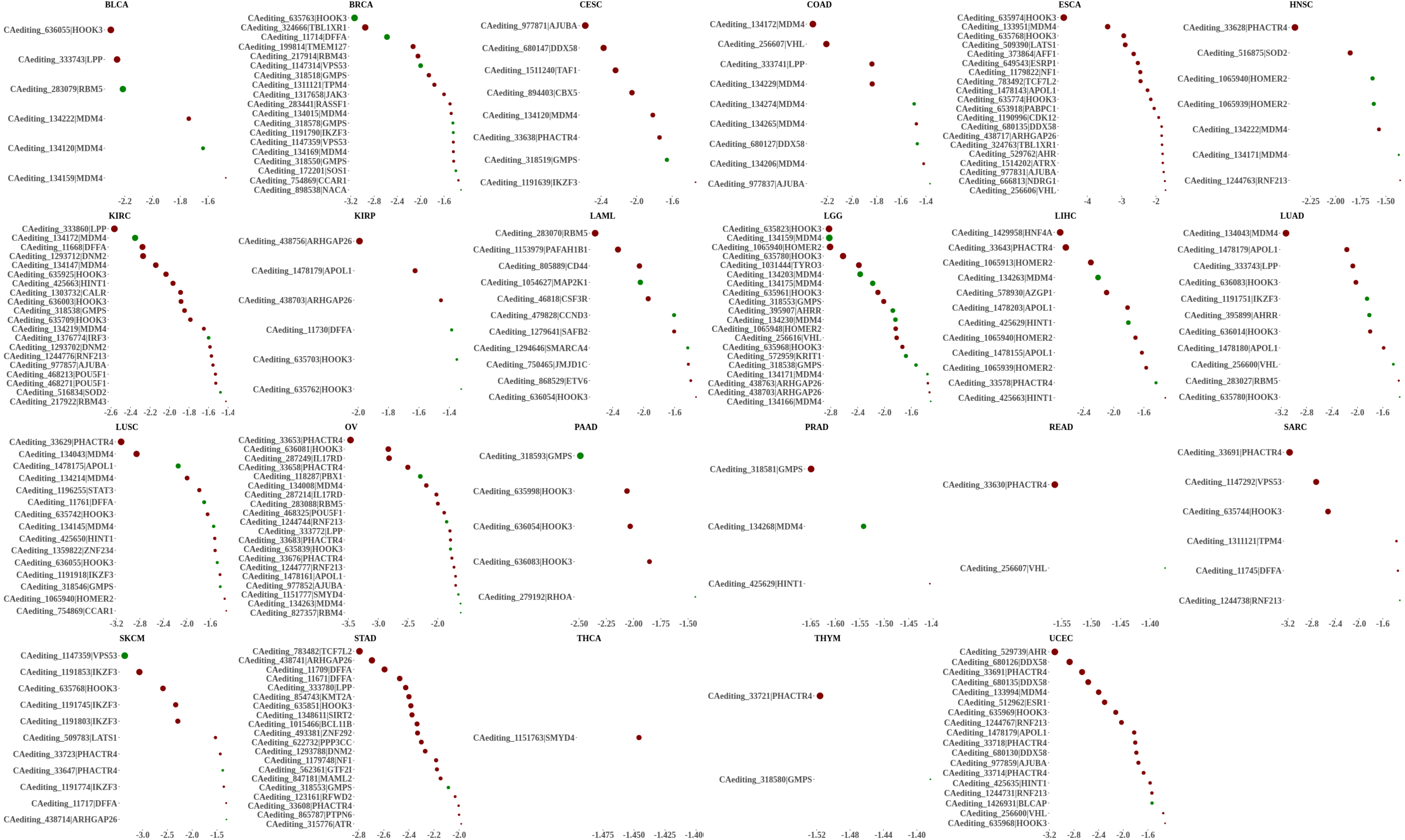

**Fig. S8. The bubble plots of top tumor survival-related RNA editing events (P < 0.05 and edited tumor samples ≧ 50) in tumor genes for each cancer type.** The x-axis shows log10(P) value, and the y-axis represents the editing events and their host tumor genes. The size of each dot denotes -log10(P) value. The red and green dots represent the RNA editing events showing high (HR > 1) or low (HR < 1) survival risks for cancer patients.

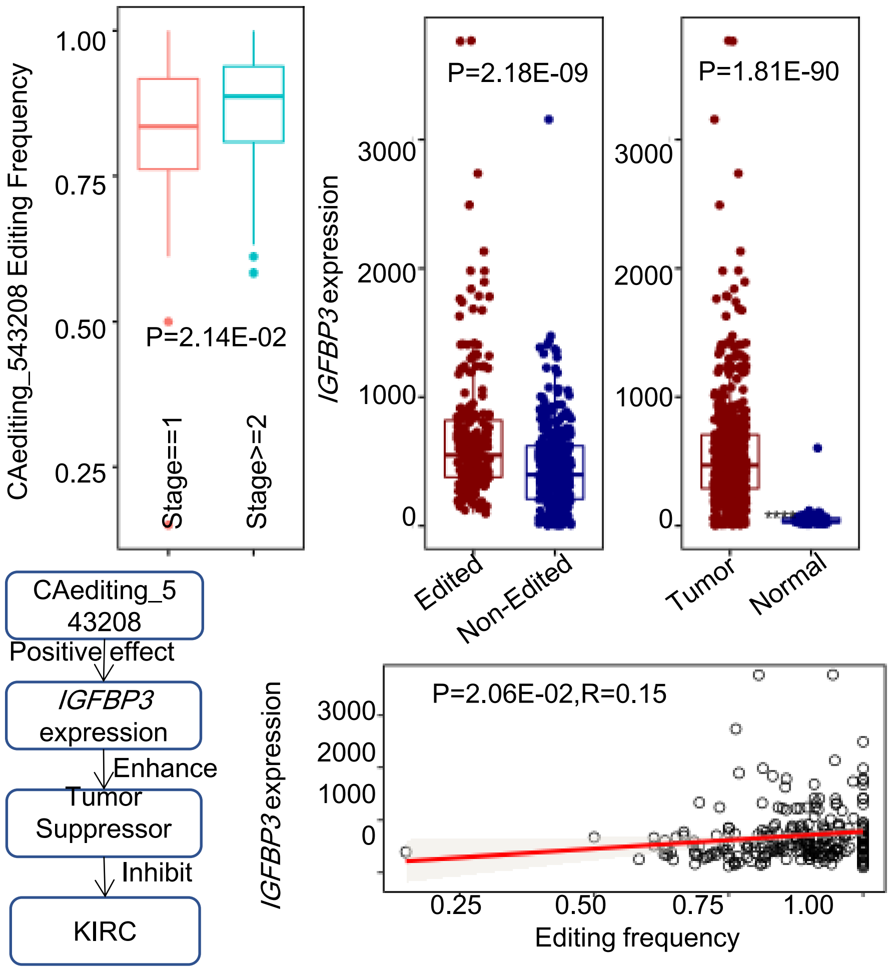


**Fig. S9. The potential of CAediting_543208 (*IGFBP3*) in KIRC cancer type.** This event occurred only in tumor samples (246/535) and none in controls (0/72), and was up-edited in tumor samples with higher stages. It was also related to the over-expressions of *IGFBP3*, which was up-regulated in tumor samples and appeared to act in an autocrine action to suppress tumor cell growth. Thus, this RNA editing event might enhance the protection functions of *IGFBP3* against cancer progression and be potential as a therapeutic target.

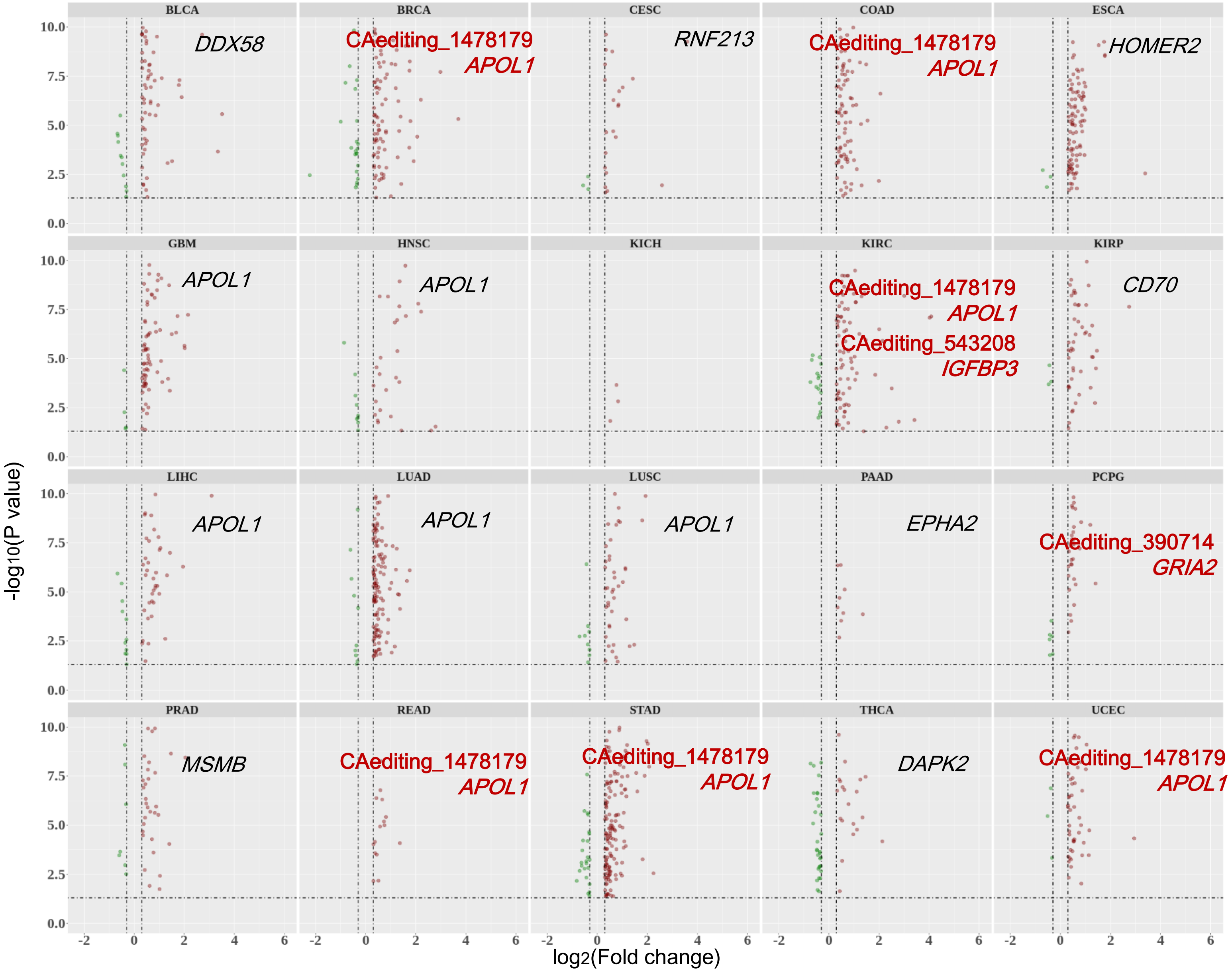


**Fig. S10. The volcano plots for DEG-associated RNA editing events in each cancer type.** These events were identified according to the procedures shown in Fig. 3E.The x-axis denotes log2(Fold change) value between RNA-edited and non-edited tumor samples, and the y-axis represents -log10(P) value. The genes shown in this figure are tumor-related genes with DEG-associated RNA editing events.

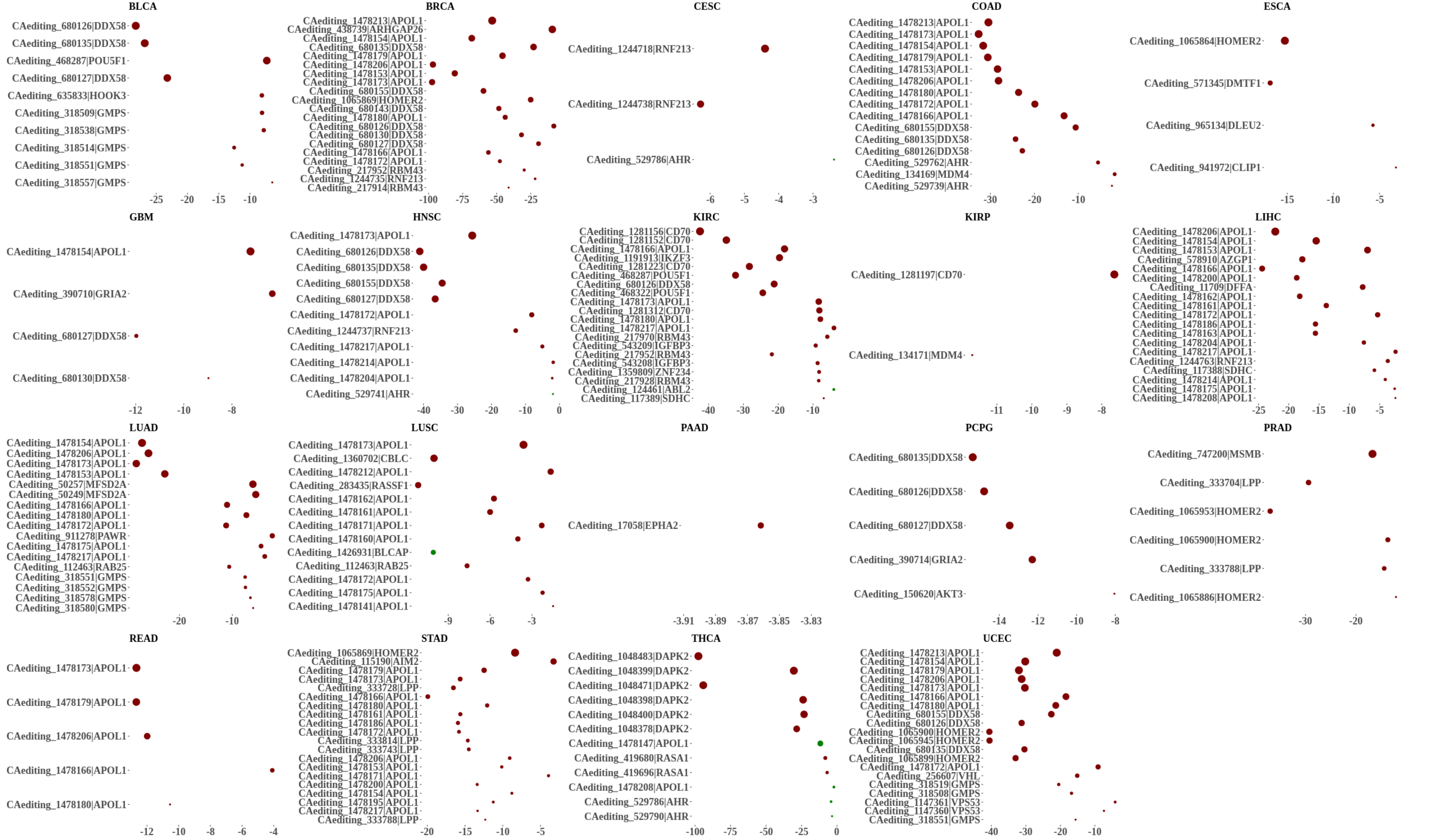


**Fig. S11. The bubble plots of top DEG-associated RNA editing events in tumor genes for each cancer type.** The x-axis shows $\log_{10}(P)$ value, and the y-axis represents the editing events and their host tumor genes. The size of each dot denotes absolute $\log_2$(Fold change) value between RNA-edited and non-edited tumor samples. The red and green dots represent the RNA editing events positively or negatively associated with the expressions of their host genes.

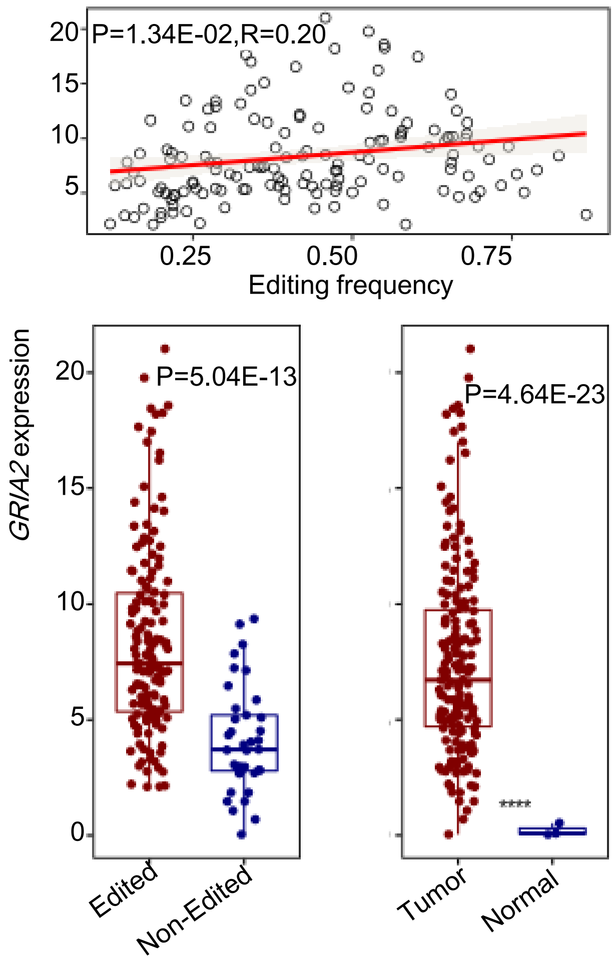


**Fig. S12. An example (chr4:157360142, CAediting_390714) showing the effects of RNA editing on gene expressions.** The frequencies of this editing event were positively associated with the expressions of its host gene (*GRIA2*) in PCPG. Since this editing event led to the significantly higher expressions of its host gene, which was also up regulated in tumor samples, CAediting_390714 may be a pathological biomarker for the PCPG cancer type. It was also supported by 146 (183) edited tumor samples and 0 (3) edited controls in this cancer type.

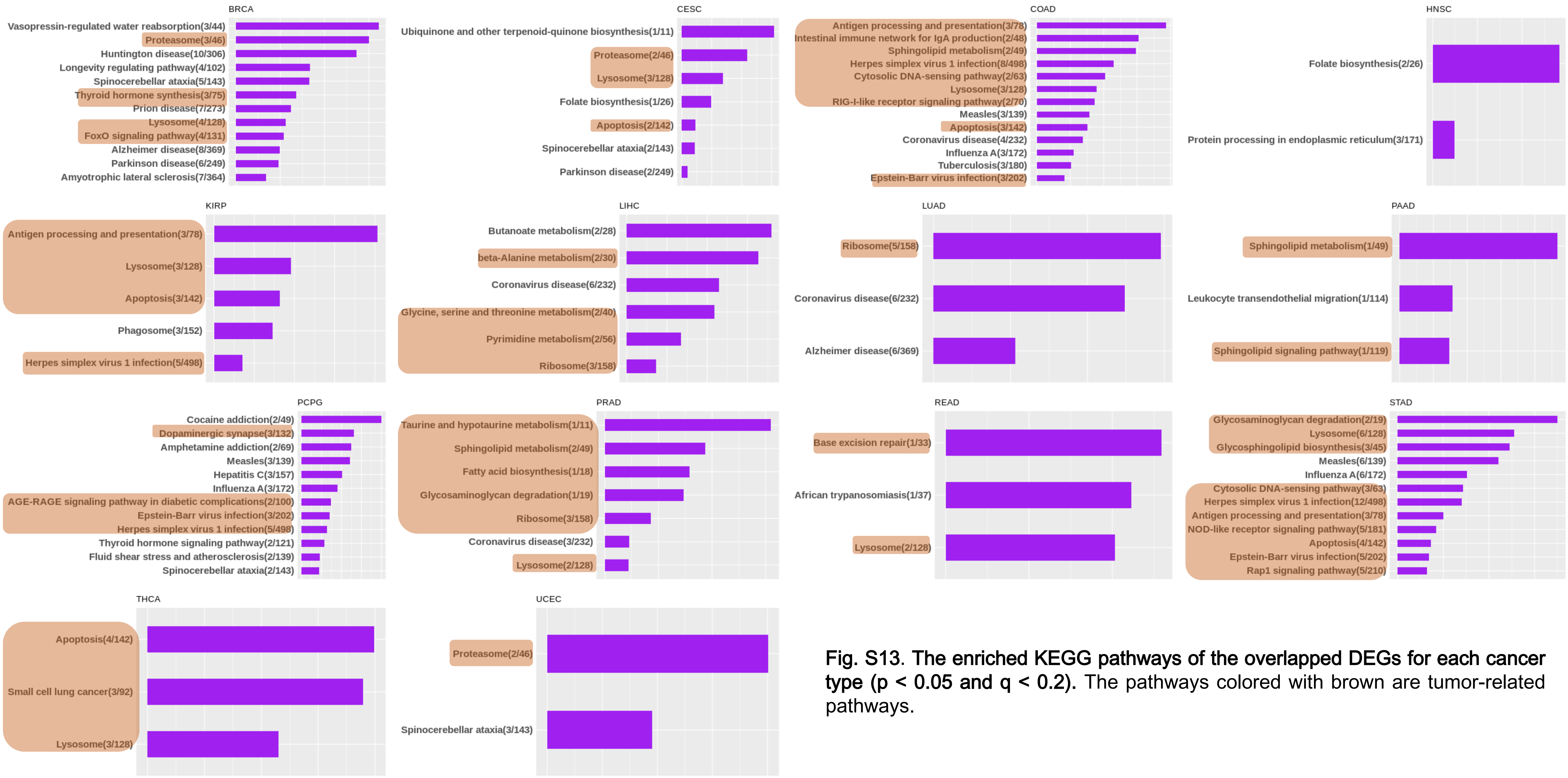


**Fig. S13. The enriched KEGG pathways of the overlapped DEGs for each cancer type ($p < 0.05$ and $q < 0.2$).** The pathways colored with brown are tumor-related pathways.

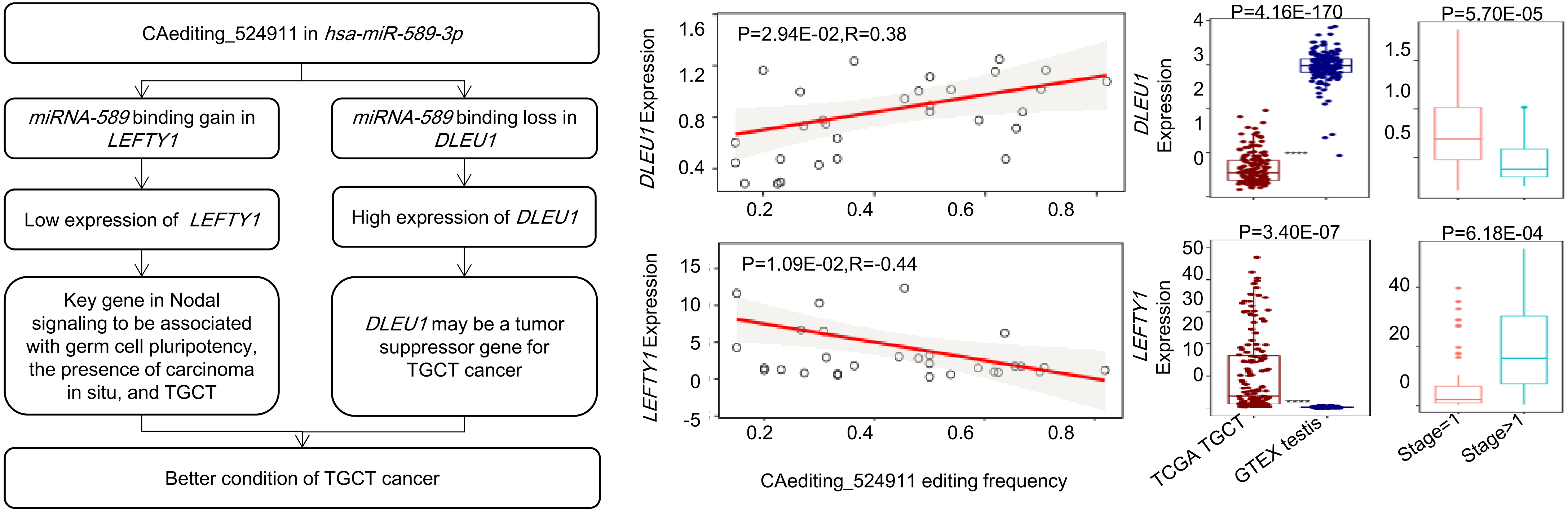


**Fig. S14. An example showing the effects of RNA editing on miRNA regulations.** An RNA editing event in chr7:5495852 (CAediting_524911) of *miR589-3p* altered the miRNA binding target from original *DLEU1* to *LEFTY1*. Due to the lost regulation of *miR-589-3p*, *DLEU1* was up-regulated in the RNA-edited TGCT tumor samples. On the other hand, *LEFTY1* was down-regulated because of the gained interactions with *miR-589-3p*.

(A) Formula $Expression_{gene(i)} = \sum_{k=1}^{N} C_k CAediting_{m(k)} + intercept$

(B) Lasso process to calculate the contributions of editing sites on

*APOL1* expressions

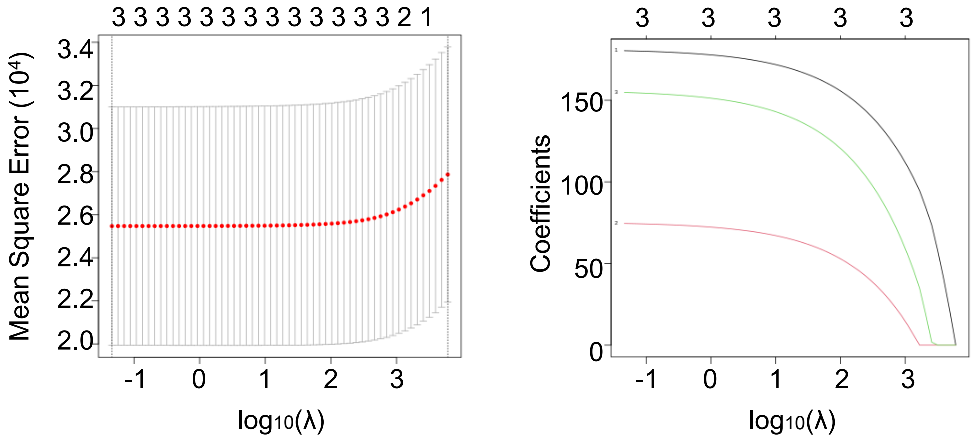


(C) Contributions coefficients of three editing sites to *APOL1* expressions

$gene = APOL1$

$CAediting = [CAediting_{1478166}, CAediting_{1478173}, CAediting_{1478179}]$

$C = [180.34, 74.55, 154.79]$

$intercept = -1.96$

**Fig. S15. An example showing the co-effects of three RNA editing events on the expressions of *APOL1* in the KIRC cancer type.** (A) The formula to estimate the co-effects of multiple RNA editing events on gene expressions. (B) The process of least absolute shrinkage and selection operator (Lasso) to calculate the contributions of three RNA editing events to *APOL1* expressions. (C) Contribution coefficients of the three editing events to *APOL1* expressions.

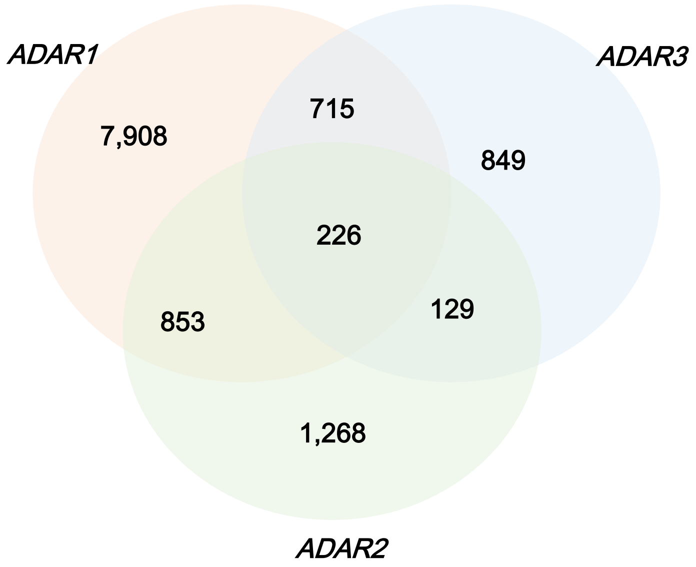


**Fig. S16. The correlations of RNA editing frequencies with *ADAR* expressions.** The frequencies of 9702, 2476, and 1919 RNA editing events are significantly associated with the expressions of *ADAR1*, *ADAR2*, and *ADAR3* respectively ($P < 0.05$ and $R > 0.3$ for Pearson method). In addition, there are also 28,062 RNA editing events ($P \geqq 0.05$ for all three *ADARs* with more than 50 edited samples) which were possibly regulated by genetic variations, splicing efficiency, RNA binding proteins, or some other mechanisms.

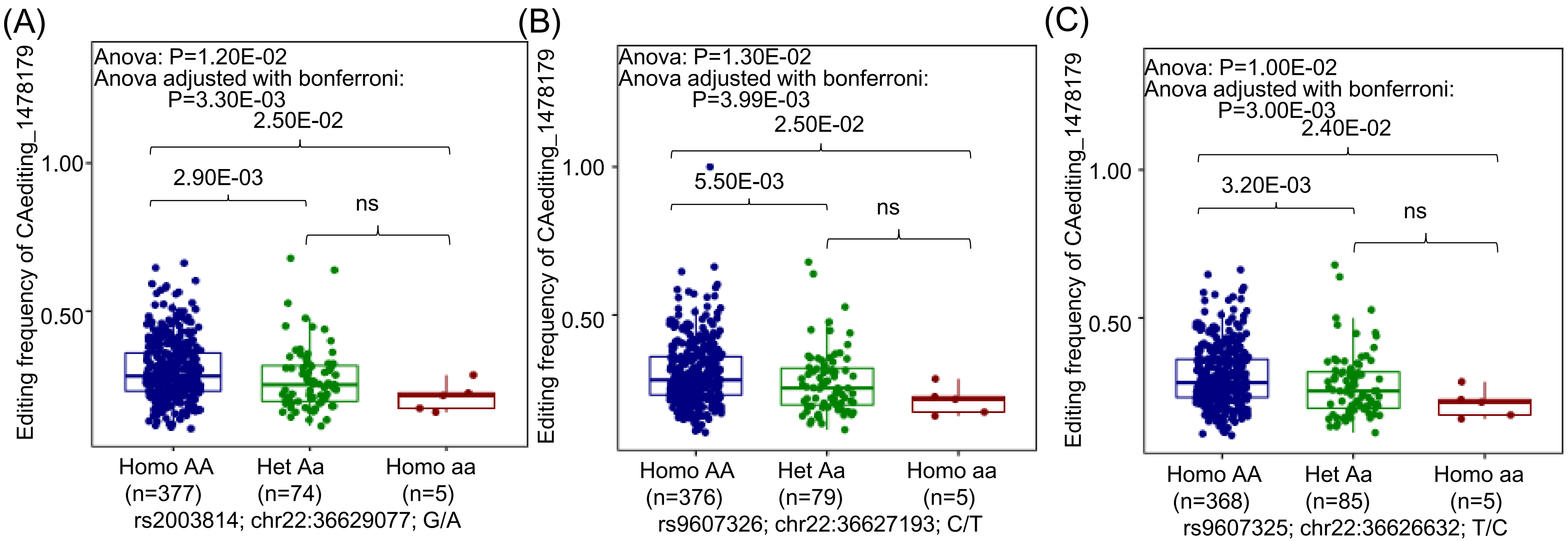


**Fig. S17. The potentially regulatory effects of rs2003814 (A), rs9607326 (B), and rs9607325 (C) on CAediting_1478179.**

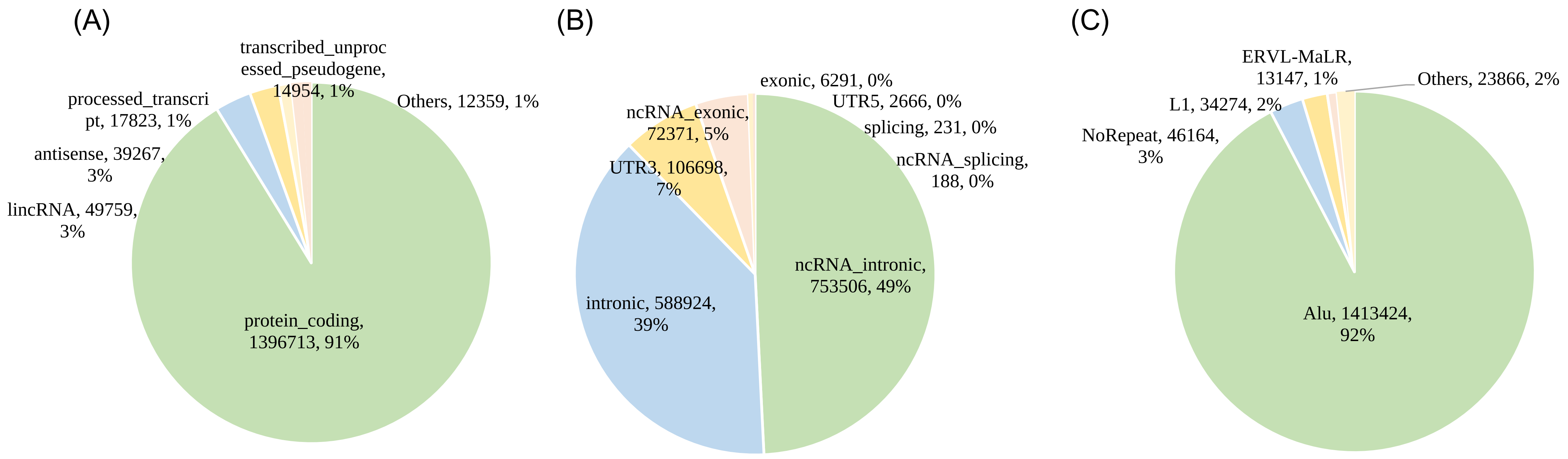


**Fig. S18. The distributions of A-to-I RNA editing events in different types of genes (A), regions (B), and repeats (C).**

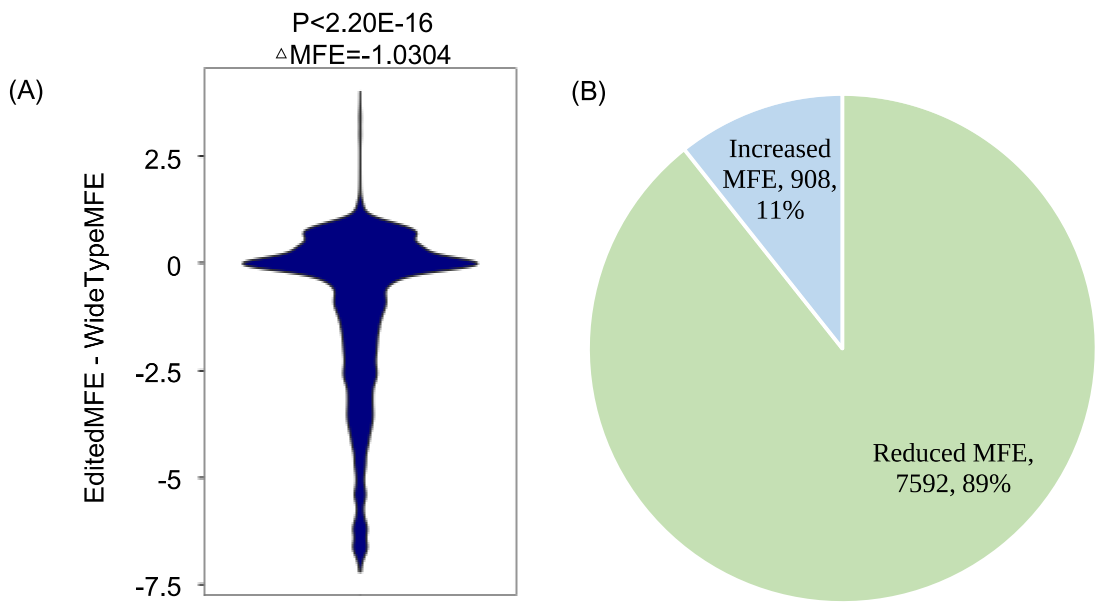


**Fig. S19. The effects of RNA editing on secondary structure.** (A) An A-to-I RNA editing event will significantly reduce the minimum free energy (MFE) to stabilize RNA structure. (B) 89.32% RNA transcripts were stabilized by one or multiple RNA editing events.

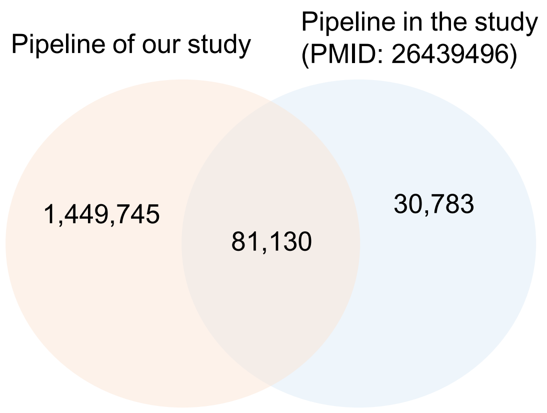


**Fig. S20. The comparisons of RNA editing events detected by different pipelines.** For comparison, we converted the genome coordinates of RNA editing events detected in previous study (PMID: 26439496) from GRCh37/hg19 to GRCh38/hg38.
